# A novel method to isolate free-floating extracellular DNA from wastewater for quantitation and metagenomic profiling of mobile genetic elements and antibiotic resistance genes

**DOI:** 10.1101/2020.05.01.072397

**Authors:** David Calderón-Franco, Mark C. M. van Loosdrecht, Thomas Abeel, David G. Weissbrodt

**Affiliations:** Department of Biotechnology, Delft University of Technology, Delft, The Netherlands; Delft Bioinformatics Lab, Delft University of Technology, Delft, The Netherlands; Infectious Disease and Microbiome Program, Broad Institute of MIT and Harvard, Cambridge, USA

**Author notes:** **Correspondence**: Prof. David Weissbrodt, Assistant Professor, Weissbrodt Group for Environmental Life Science Engineering, Environmental Biotechnology Section, Department of Biotechnology, Faculty of Applied Sciences, TU Delft, van der Maasweg 9, Building 58, 2629 HZ Delft, the Netherlands.

**Keywords:** Free-floating extracellular DNA, Intracellular DNA, Wastewater samples, Xenogenetic Elements, Antibiotic resistance, qPCR, Metagenomics

## Abstract

Antibiotic resistant genes (ARGs) and mobile genetic elements (MGEs) can be found in the free-floating extracellular DNA (eDNA) fraction of microbial systems. These xenogenic components can generate bacterial cells resistant to one or more antibiotics by natural transformation. Because of low concentration in wastewater, the obtaining of a high quality and a high yield of eDNA extract is challenging. We developed a method using chromatography to isolate eDNA without causing cell lysis (often unchecked) from complex wastewater matrices. The chromatographic step involved a diethylaminoethyl-cellulose-monolithic column to capture the eDNA found in cell-free filtered wastewater samples (e.g. influent wastewater, activated sludge and treated effluent wastewaster). Free-floating eDNA yields from 1 L of influent, activated sludge and treated effluent water reached 12.5 ± 1.9 μg, 12.3 ± 1 μg and 5.6 ± 2.9 μg of raw eDNA and 9.0 ± 0.7 μg, 5.6 ± 0.46 μg and 2.6 ± 1.3 μg of purified eDNA, respectively. In order to check the suitability of free-floating eDNA extracts for molecular analysis, qPCR and metagenomics were performed. eDNA extracts from treated effluent water were analyzed by qPCR to quantify a selected panel of ARGs and MGEs. Microbiome, resistome, and mobilome profiles from activated sludge free-floating eDNA were measured by metagenomic sequencing. Between iDNA and eDNA fractions, qPCR showed differences of 0.94, 1.11, 1.92 and 1.32 log_10_ gene copies mL^−1^ for sulfonamides resistant genes (*sul1* and *sul2*), β-lactamase resistance gene *bla_CTXM_*, and the class 1 integron-integrase (*intI1*) MGE, respectively. These differences highlighted the crucial need for an isolation method to discern both iDNA and eDNA to understand ARGs persistence and quantity in complex cultures. The eDNA yields obtained from 1 L of activated sludge (3.6 g of total suspended solids L^−1^) samples were substantially higher than the amount of DNA template needed for high-throughput sequencing (>1 μg) in service facilities. Subsystems classification showed that the eDNA metagenome was mainly composed by MGEs (65.1%). The 35.9% rest related to traditional functional genetic signatures. It was the first time the resistome from the eDNA fraction was analyzed showing lower number of primary aligned reads when compared to the iDNA and a predominance of aminoglycosides and β-lactamams. Metagenome results showed that eDNA can not be discarded as a pool of ARGs and MGEs for horizontal gene transfer. This novel isolation method was powerful to elucidate the molecular compositions of free-floating eDNA fractions in complex environmental samples such as wastewater environments at different microbial densities. Data obtained using this extraction method will foster xenogenic and microbial risk assessments across urban and natural water systems. This will support water authorities in the delineation of measures to adopt at wastewater treatment plants to remove them and safeguard environmental and public health.

**Graphical abstract:** Picture created with BioRender

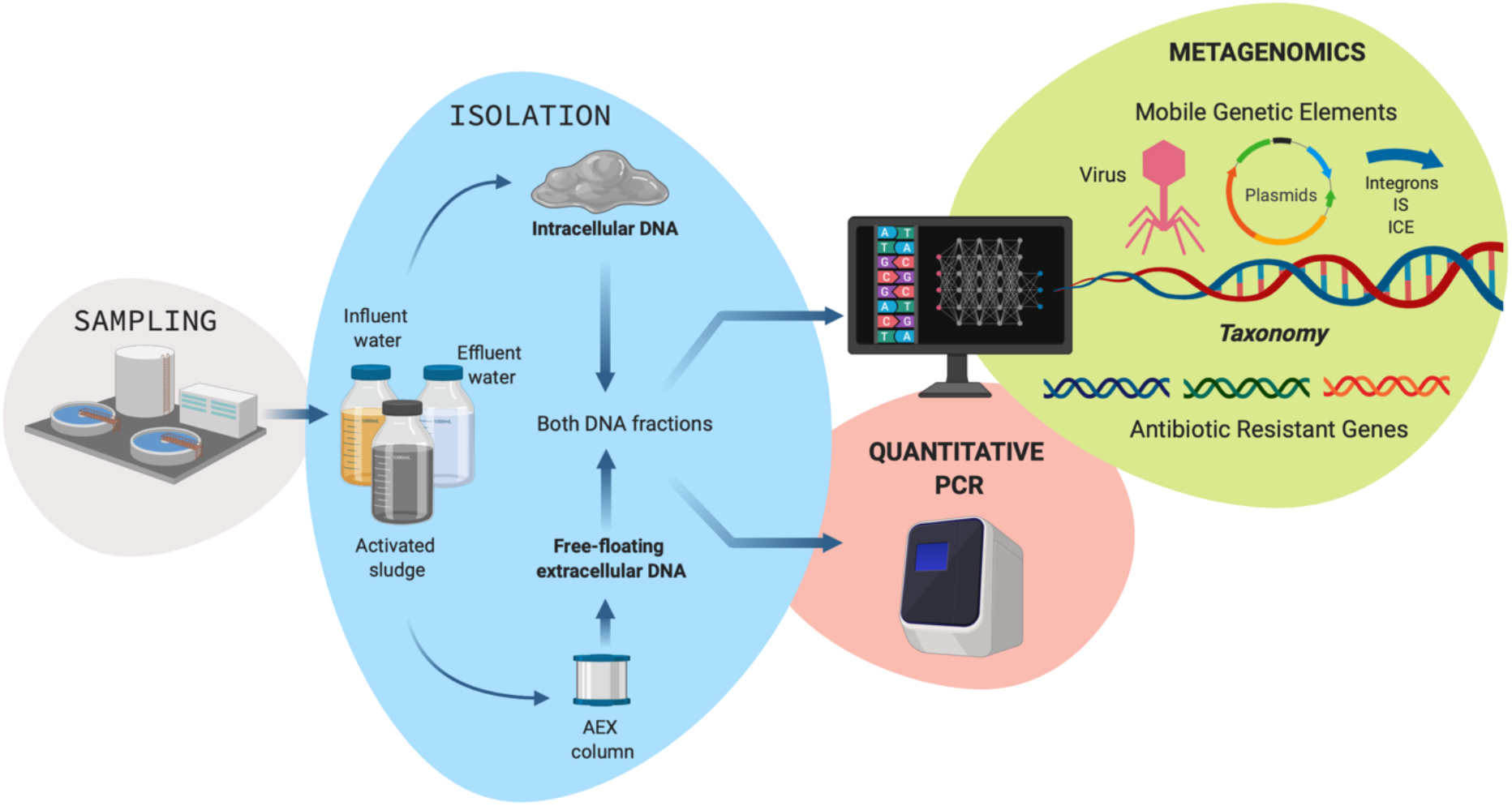

## Introduction

Extracellular DNA retrieved from environmental samples can be defined as the DNA fraction that is strictly not embedded inside cell membranes^1^. In contrast, DNA enclosed inside microorganisms is called intracellular DNA (iDNA). eDNA can be found in various environments *e.g.*, water systems ^1–3^, soil ^4^, animal and human guts and fluids^5^. The acronym “eDNA” is widely intermixed to describe either environmental DNA or extracellular DNA. Environmental DNA does not necessary discriminate between extracellular and intracellular fractions of nucleic acids. It only means that DNA extracted originates from cells that were present in the investigated ecosystem at the sampling event ^6^. In such studies, environmental DNA has normally been extracted from biological samples that have been concentrated on filtration columns ^7^ or membrane filters that collects all kinds of DNA^8^.

Extracellular DNA (hereafter referred to as eDNA) is abundant and plays an important role as structural component of microcolonies. At meso scale, it contributes in stabilizing microbial bioaggregates from pure-culture to multi-species flocs ^9^, granules ^10,11^, and biofilms engineered for wastewater treatment. Hence, eDNA is a key component within the matrix of extracellular polymeric substances ^9,12–14^. Whether eDNA is actively or passively released in these samples primarily remains a mystery, although some authors provide hypothetical claims about one or the other art ^15,16^.

Active or passive release out of cells are the main sources of eDNA ^1,17^. Active release comprises all cellular mechanisms that are involved by microorganisms to extrude DNA from their cytoplasm ^18,19^. Passive release consists in the release of intracellular DNA from cell lysis, either necropsied or infected by a virus^17^. Both active and passive releases of DNA can contribute to biofilm formation. Until mechanistic measurements will be made available, there will still be strong debate on the ‘intention’ of cells to release DNA to drive biofilm formation.

We define free-floating extracellular DNA as “all the DNA components that are neither enclosed inside cells nor in complex matrices and that are persistingly floating in aqueous samples”. Free-floating DNA is a concept normally associated with environmental DNA and fish ecology and diversity ^20^. Thanks to its ability to be adsorbed on sediments, eDNA can represent a genetic proxy of microbial and cellular diversity among different biogeographical areas ^1,21^. Isolating and analyzing eDNA in environmental samples provides insight on the dynamics, interactions and evolutionary history of populations of microorganisms and higher organisms that are or have been present in the investigated environment. The persistence of eDNA on biofilms and of free-floating eDNA sorbed or not on different surfaces may generate hotspots for horizontal gene transfer (HGT) in microbial biocoenoses. Natural competence is a widely distributed cellular mechanisms harboured by microorganisms in nature to take up molecular resource from their surroundings. Naturally competent microbes may take up uptake free or bound eDNA ^17^. Natural transformation is a parasexual mechanism for exchange of genetic material induced by stress conditions such as nutrient limitations or the presence of antibiotics ^22^. Such conditions are found in densely populated cultures such as activated sludge samples. Natural transformation in managed environments does not implicitly result in a threat for human health, since it is a mechanisms for generating diversity and adaptation ^23^.

However, if antibiotic resistant genes, mobile genetic elements, and pathogenic islands can be found in eDNA fractions, the formation of pathogenic bacterial cells resistant to one or more antibiotics, also known as superbugs can be induced. It has already been shown that DNA release by sterilized cultures with conventional industrial and research facilities methods does not lose its integrity and capacity to be re-amplified. Thus, making it a source for GMOs DNA fragments as well as ARGs and MGEs ^24^. This unfavorable consequence is undesired. Mechanisms of superbugs generation and the underlying sources of genetic material that can prompt it need to be studied. Highly effective methods to obtain eDNA from complex biological samples are needed in order to obtain DNA templates of high quality in high quantity for quantitative and metagenomics studies.

In the last years, some methods to extract eDNA have been published. As examples, Wang *et al.* (2016) have used an aluminum hydroxide column to adsorb aquatic eDNA. Nagler *et al.* (2018) have opted for sequential steps involving enzymatic treatments to extract eDNA from cattle rumen samples. Yuan et al. (2019) have involved magnetic beads in combination with the classic DNA precipitation method involving the surfactant cetyl trimethyl ammonium bromide (CTAB), yielding good amounts of eDNA (2.3 μg out of 5 mL wastewater sample). Silica solid phases, similar to commercial silica resin columns, have been tested to adsorb and extract eDNA from low-concentration clinical samples ^27^. Cell lysis has seldom been investigated (although often debated) across protocols for eDNA extractions. Assessing and preventing cell lysis during eDNA isolations is crucial to obtain confident analytical results from eDNA templates. This is important to characterize and compare the molecular compositions of differ3ent DNA fractions at high resolution.

Here, we present for the first time a novel reusable chromatographic method to isolate free-floating eDNA at high quality and yield from complex wastewater matrices. It requires relatively small sample volumes (1000 mL). Cell lysis during the extraction steps is evaluated. Overall, isolated eDNA samples are commercially sufficient for being processed for metagenomics and quantitative studies.

## Material and Methods

### Sampling from the influent, activated sludge tank and effluent of a wastewater treatment plant

Biological samples were collected from the urban wastewater treatment plant (WWTP) Harnaschpolder (Waterboard Delfland, The Netherlands) operated for full biological nutrient removal. Samples of influent and effluent water were also collected from the same installation.

Influent water was collected as the inlet after primary treatment of the WWTP Harnashpolder. Three biological replicated were collected in three different days. A total of 1000 mL of influent water per replicate was collected. All samples were processed in a timeframe of less than 2 h prior to DNA extraction.

Six biological replicates of activated sludge were collected in two sampling campaigns as grab samples from the activated sludge tank. Each campaign consisted of three successive dry days, *i.e.,* without recent rainfall and variations in hydraulic retention time. All raw activated sludge samples were stored at 4 °C in a timeframe of less than 2 h prior to isolations of free-floating eDNA and extractions of iDNA. A total volume of 1000 mL of activated sludge was collected per replicate.

Effluent water was collected at the outlet of the tertiary treatment of WWTP Harnashpolder. Three biological replicates were collected in three different days. A total of 1000 mL of treated water per replicate was collected. All samples were processed in a timeframe of less than 2 h prior to DNA extraction.

### Isolation of free-floating extracellular DNA from activated sludge and effluent water samples

The filtered supernatants were loaded on a positively charged 1-mL diethylaminoethyl cellulose (DEAE) chromatographic column (BIA Separations, Slovenia). The column was preminarily equilibrated at a flowrate of equilibration buffer of 0.6 mL min^−1^ and maintaining the pressure below the maximum limit of 1.8 MPa. Because of very high porosity, reusability amd flow characteristics (up to 16 mL min^−1^), this monolithic chromatographic column is is an efficient tool for separation or purification of large biomolecules such as genomic and viral DNAs ^28^. A schematic representation of the process is sketched in **figure 1**.

**Figure 1.**
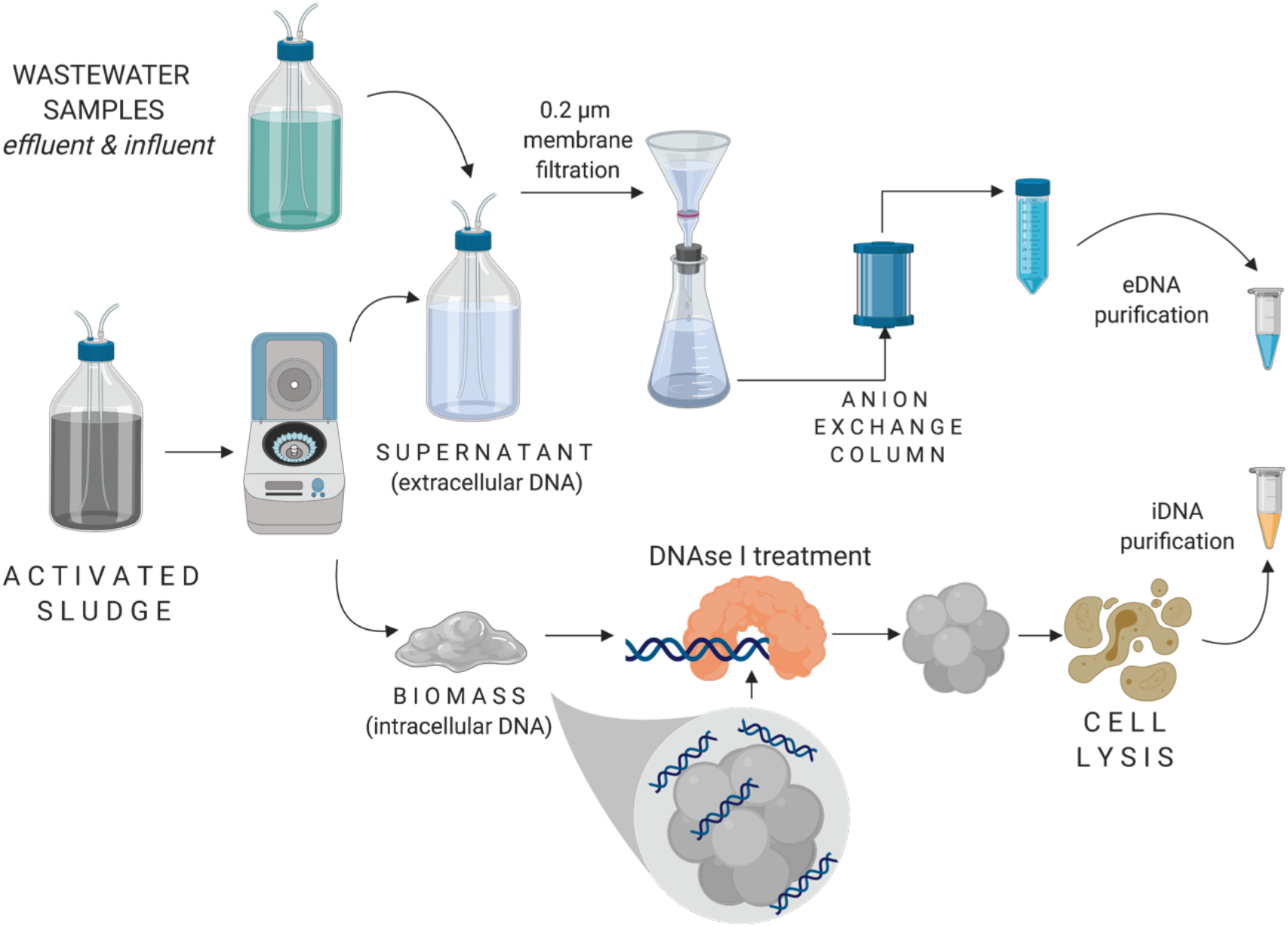
Schematic representation of the free-floating eDNA and iDNA isolation method. *Picture created with BioRender*

Buffers and solutions were used to equilibrate, elute, regenerate, clean, and store the column. The equilibration buffer consisted of a mixture at pH 7.2 of 50 mmol L^−1^ Tris and 10 mmol L^−1^ EDTA. The elution buffer was a mixture at pH 7.2 of 50 mmol L^−1^ Tris, 10 mmol L^−1^ EDTA, 1.5 mol L^−1^ NaCl. The regeneration buffer was a mixture at pH 7.2 of 50 mmol L^−1^ Tris, 10 mmol L^−1^ EDTA, 2 mol L^−1^ NaCl. The cleaning solution comprised 1 mol L^−1^ NaOH and 2 mol L^−1^ NaCl. The storage solution consisted of 20% ethanol in ultrapure water (Sigma Aldrich, USA). Column preparation and processing were performed according to manufacturer’s instructions. The elution of eDNA was tracked over time using an HPLC photodiode array detector (Waters Corporation, USA) recording the UV-VIS absorbance at the absorbance wavelength characteristic of DNA (260 nm). The eluted eDNA fraction was further treated sequentially with absolute ethanol and a solution of 70% ethanol in ultrapure water (Sigma Aldrich, USA) ^29^ to precipitate the raw eDNA. The precipitated raw eDNA was incubated with proteinase K (Sigma-Aldrich, UK) at 0.85 g L^−1^ during 2 h in order to digest remaining co-extracted proteins (*e.g.*, DNA-bound proteins). The enzymatic reaction was stopped in a heat block at 50°C for 10 minutes. The precipitated and protein-digested raw eDNA extract was finally purified using a GeneJET NGS Cleanup Kit (Thermo Scientific, USA). The purified eDNA isolates were stored at −20°C pending molecular analysis.

### Extraction of intracellular DNA from biomass separated from activated sludge and effluent water

Biomass was obtained as pellet from the centrifugation step (activated sludge) and filtration step (effluent water). An amount of 0.25 g (3.6 g TSS L^−1^) activated sludge was processed to extract intracellular DNA using a NucleoSpin® Soil kit (Macherey-Nagel) according to the manufacturer’s instructions, after a preliminary incubation of 1 h with 300 U mL^−1^ DNase I to remove the remaining eDNA from the sample.

Biomass from effluent water was processed as follows. A volume of 500 mL of effluent was filtered through a 0.22 μm PVDF membrane (Pall corporation, USA). Filters were frozen at −20°C until extraction. The intracellular DNA present on filters were extracted with the DNeasy kit Power Water (Qiagen, NL) following manufacturer’s instructions.

DNA quality and quantity were analyzed using NanoDrop spectrophotometer (ND-1000, USA) and Qubit® dsDNA assays (Thermo Fisher Scientific, USA), respectively.

### Plasmid DNA used as chromatography selectivity control

A bacillus expression pHT01 plasmid (MoBiTec GmbH, Germany) was used as selectivity control for the DEAE column.

### Biomass pre-treatment, live-dead staining, and flow cytometry analysis

Flow cytometry was used to test for cell lysis in combination with live-dead staining. In order to disaggregate cells (a prerequisite for migration of individual cells in the flow cytometer), activated sludge samples were firstly diluted at 1:500 in 1x PBS buffer. The diluted sludge samples were then mild-sonicated (Branson Sonifier 250, USA) on ice in 3 cycles of 45 s at 40 W. After sonication, samples were diluted at 1:500 in 1x PBS buffer. Right after the second dilution, samples were filtered through a 10 μm syringe filter in order to remove any remaining cell debris and membranes.

The live-dead staining protocol and flow cytometry analyses were based on previous works ^30–33^. Two staining dyes were used to track viable cells with SYBR green (Invitrogen, USA) and dead cells with red-fluorescent propidium iodide(PI) (Invitrogen, USA). Two working solutions were prepared from 10’000x SYBR Green and 30 mmol L^−1^ PI dissolved in di-methyl sulfoxide (DMSO) stock solutions.

To obtain total stained cell count, 10 μl of 10000x SYBR green stock solution was diluted in TRIS-HCl (pH 8), obtaining a final 100x SYBR green working solution. To obtain the viable cell count, a final working solution of 100x SYBR green and 6 μmol L^−1^ PI was used. Working solutions were thoroughly mixed by homogenization using a vortex and stored at −20°C in the dark pending analysis.

Volumes of 5 μL of each of the staining working solutions was added to 495 μL of pre-treated biomass sample. After stains addition, samples were incubated at 37 °C for 10 minutes. After incubation, samples were kept in the dark until flow cytometry measurement.

Flow cytometry measurements were performed in a BD Accuri C6 flowy cytometer (BD Biosciences, Belgium) equipped with a 50 mW laser exciting analytes at a fixed wavelength of 488 nm. Fluorescence intensity was collected at emission wavelengths of the green (FL1 = 533 ± 30 nm) and the red (FL3 > 670 nm) fluorescence detection channels. Measurements were performed at a flow rate of 66 mL min^−1^ on 50 μL sample volume with a threshold value of 700 on FL1 to reduce the background detection noise.

### Gel electrophoresis on the extracellular DNA

eDNA samples were analyzed by gel electrophoresis with agarose at 1% (w/v) (Sigma-Aldrich, Haverhill, United Kingdom) in 1xTAE buffer.

### Quantitative polymerase chain reaction (qPCR) analysis of selected ARGs and MGEs on eDNA and iDNA from effluent water samples

A panel of genes was selected for qPCR analysis on eDNA and iDNA fractions extracted from effluent water samples. The 16S rRNA gene was selected as a proxy to quantify total bacteria. ARGs and MGEs were selected from a panel used for wastewater samples ^34^. Standards for qPCR were generated from ResFinder (https://cge.cbs.dtu.dk/services/ResFinder/), a curated database of ARGs. The chosen ARGs confer resistance to antibiotics with the highest consumption in The Netherlands, namely: macrolides (multidrug export protein gene *ermB*), sulfonamides (sulfonaminde resistance genes *sul1* and *sul2*), fluoroquinolones (quinolone resistance gene *qnrS*) and extended-spectrum β-lactamase (cefotaxime-hydrolyzing β-lactamase *bla_CTXM_*) **(Table 1)**. The class I Integron-integrase gene *intI1*, known to be responsible of some horizontal gene transfer phenomena ^35^ was included to assess the presence of MGE. Standards, primers and reaction conditions used in this study are listed in the supplementary material and **tables S1** and **S2** in supplementary matrial.

**Table 1.**
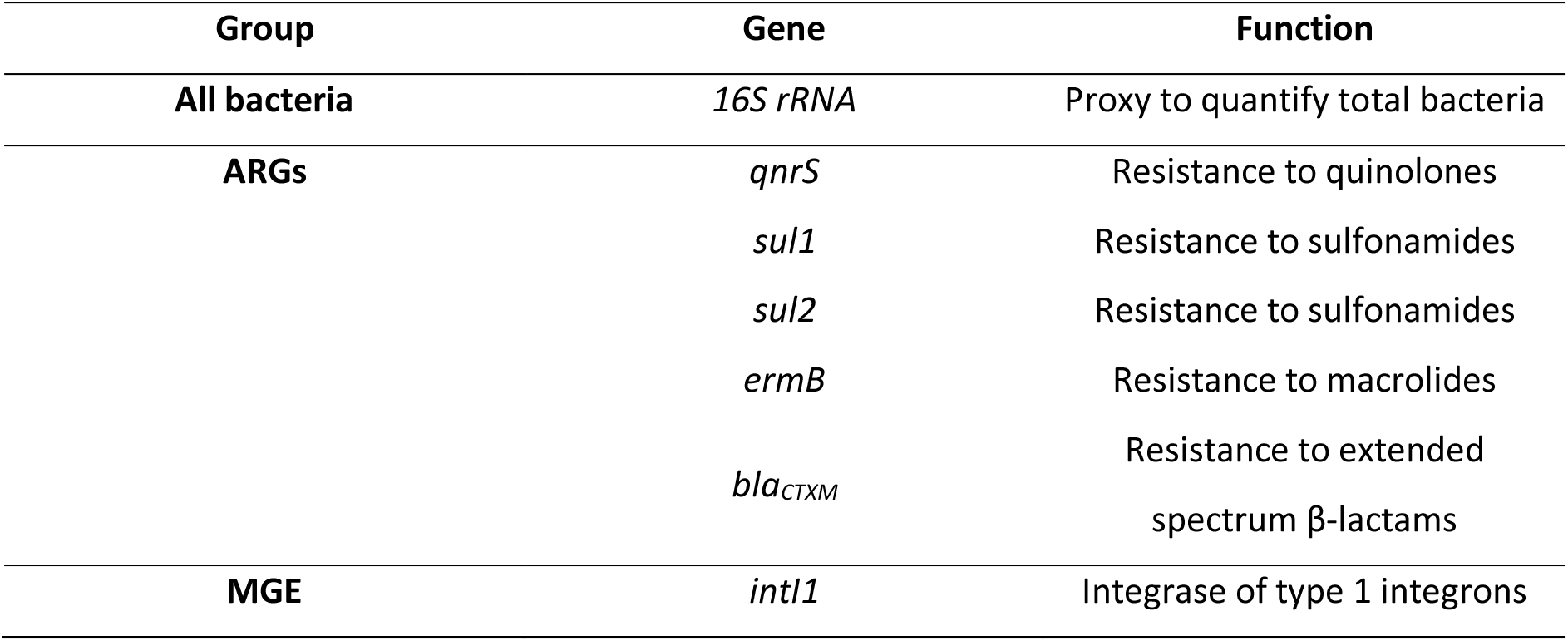
Genes analysed by qPCR on eDNA and iDNA fractions obtained from activated sludge and effluent water samples. Genes are classified in three groups of interest: all bacteria (16S rRNA gene), antibiotic resistance genes (ARGs), and mobile genetic element (MGE).

### Metagenomics profiling of eDNA and iDNA fractions from activated sludge samples

Libraries of eDNA and iDNA samples at 50 ng μL^−1^ were sequenced using a MiSeq PE300 benchtop sequencer (Illumina, USA). Library preparation was done with a TruSeq DNA PCR-Free kit (LT Lib PREP KT-ST B PhiX control v3). Datasets of 14 M reads were obtained per sample corresponding to a sequencing depth of 6.5 GB for iDNA and 6.8 GB foreDNA fractions.

The quality of the aquired Illumina reads was assessed by FastQC version 0.11.9 with default parameters^36^, open-source software and paired-ends reads were trimmed and filtered by Trimmomatic version 0.39 with default parameters ^37^. Alignments were performed by BWA-mem version 2 with default parameters ^38^, generating a SAM file. To filter soft and hard clipped reads, SAM files were filtered by Samclip tool with default parameters, removing undesirable alignments that could generate downstream problems (https://github.com/tseemann/samclip). For taxonomic microbial classification, Centrifuge^39^ was applied (http://ccb.jhu.edu/software/centrifuge). To identify antibiotic resistance genes, we used the MEGARes database (http://megares.meglab.org). The metagenomes were also searched for signatures of known mobile genetic elements through the ACLAME (http://aclame.ulb.ac.be) for plasmids, prophages and viruses ^40^. The mobilome was also searched for integrative conjugative elements through the ICEberg database (http://db-mml.sjtu.edu.cn/ICEberg/) ^41^, insertion sequences through ISfinder (https://isfinder.biotoul.fr) ^42^ and integrons through the INTEGRALL database (https://integrall.bio.ua.pt) ^43^.

Both MEGARes and ACLAME were used as reference where trimmed and filtered metagenomic reads were aligned to. Output SAM file from BWA-mem alignment was converted into a BAM file using SAMtools version v1.4 ^44^ available in (https://github.com/samtools/samtools). We removed unmapped reads and ignore secondary alignments to get a list of the best hits for further data processing. ARGs and MGEs hits were considered when more 2 or more reads per variant were aligned.

### Comparative subsystems analysis of the metagenomic data

Unassembled clean reads from activated sludge eDNA and iDNA fractions were annotated by Metagenome Rapid Annotation using Subsystem Technology (MG-RAST) server (http://metagenomics.nmpdr.org/), an open-access metagenome curation and analysis platform ^45^.

### Statistics

Statistical analyses were performed on all molecular datasets with R 3.5.1 ^46^ and RStudio (https://www.rstudio.com/). For the analysis of significance on the purification effect on the yield of DNA purification, a non-paremetric Wilcoxon test was performed.

A parametric two-tailed Student’s t-test with statistical significance established at the 95% confidence level (p<0.05) was performed to analyze the significance of the differences in ARG and MGE compositions between eDNA and iDNA fractions.

## Results and Discussion

### High yields of free-floating eDNA were obtained by chromatographic isolation

The diethylaminoethyl cellulose (DEAE) column was efficient to isolate and concentrate the free-floating eDNA fraction from large volumes of influent wastewater, activated sludge, and effluent water of the WWTP. The column performed with high nucleic acid selectivity for the retention, separation, and spectrophotometric detection (λ = 260 nm) with known concentration of pure pHT01 plasmid of *Bacillus subtilis* as control (**Figure S1a**). The raw non-precipitated eDNA obtained from filtered activated sludge started to elute after 10 minutes for 35 minutes (**Figure S1b**). The mode of the chromatographic peak was detected at a retention time of 18 min. The concentrated raw free-floating eDNA extract displayed a hydrogel aspect (**Figure 2**). Its color related to the source of the eDNA extract, namely dark brown with influent water, yellow with activated sludge or colorless with effluent water. The explanation could relate to humic acids. Humic acids are abundant in wastewater and can interact with DNA ^47^. They adsorb onto activated sludge ^48^ while effluent water is almost deprived of them ^49^.

**Figure 2.**
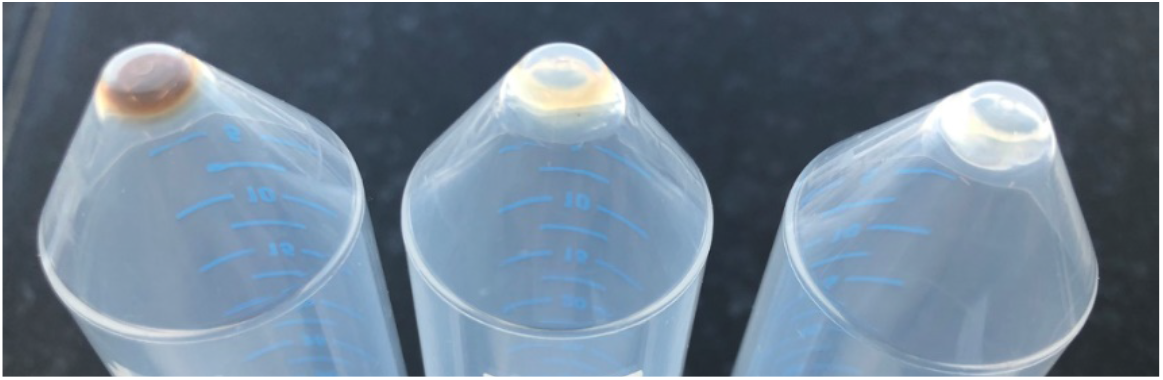
Aspects of raw extracellular DNA extracts obtained from different wastewater environments. From left to right: (i) filtered influent water (brown free-floating eDNA extract due to high presence of humic acids), (ii) centrifuged and filtered activated sludge supernatant (yellow extract), and (iii) filtered effluent water (colorless extract).

The yields (*i.e.*, mass of DNA isolated in molecular biology) of raw and purified free-floating eDNA fractions achieved from influent wastewater, activated sludge and effluent water are shown in **Figure 3**. Yield and concentrations are summarized in **Table 2**. Only eDNA obtained from activated sludge samples showed significant differences in the yields between raw and purified eDNA (p<0.05). There is no significant difference (p>0.05) between raw eDNA obtained from different water quality samples. The influent wastewater displayed a significantly (p < 0.05) higher fraction of purified free-floating eDNA (9’000 ng) than activated sludge (5’631 ng) and effluent water (4’276 ng) even if this may be due to human factors and extraction kit used.

**Table 2.**
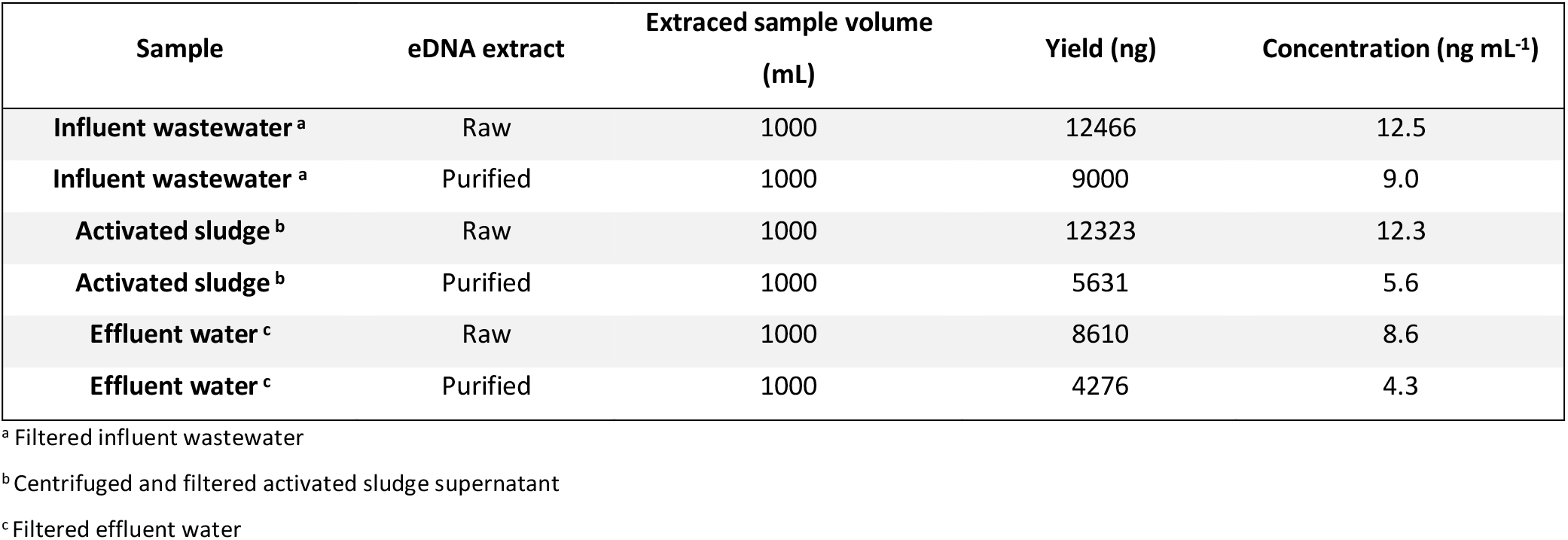
eDNA yield and concentrations for different water qualities before and after commercial kit purification. Extracting volume corresponds to the volume of initial water sample from which eDNA had been extracted.

| Sample | eDNA extract | Extraced sample volume (mL) | Yield (ng) | Concentration (ng mL<sup>-1</sup>) |
| --- | --- | --- | --- | --- |
| Influent wastewater <sup>a</sup> | Raw | 1000 | 12466 | 12.5 |
| Influent wastewater <sup>a</sup> | Purified | 1000 | 9000 | 9.0 |
| Activated sludge <sup>b</sup> | Raw | 1000 | 12323 | 12.3 |
| Activated sludge <sup>b</sup> | Purified | 1000 | 5631 | 5.6 |
| Effluent water <sup>c</sup> | Raw | 1000 | 8610 | 8.6 |
| Effluent water <sup>c</sup> | Purified | 1000 | 4276 | 4.3 |

<sup>a</sup> Filtered influent wastewater
<sup>b</sup> Centrifuged and filtered activated sludge supernatant
<sup>c</sup> Filtered effluent water

**Figure 3.**
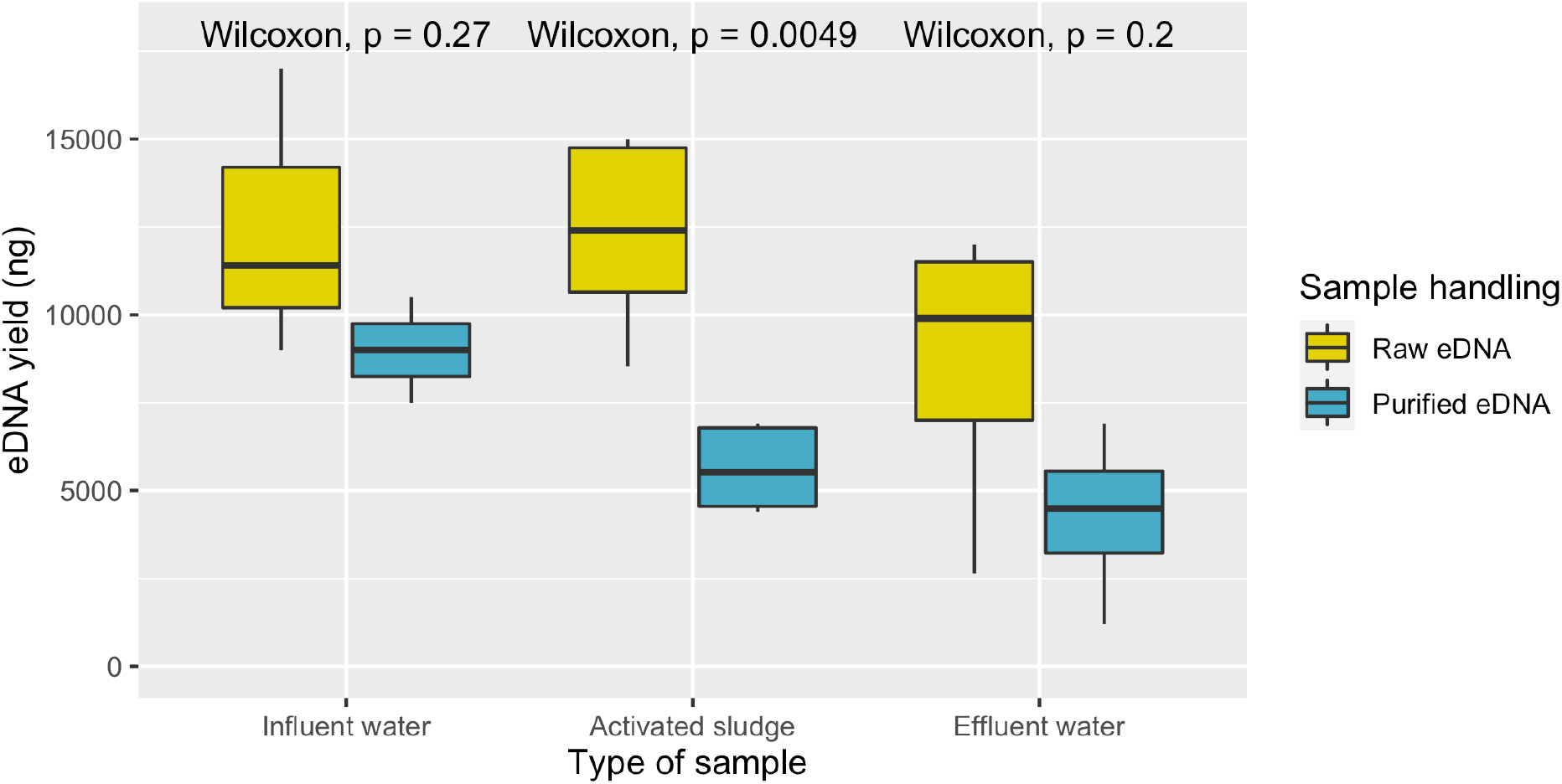
Effect of purification on eDNA yields depending on different water qualities: influent water, activated sludge and effluent treated water.

The raw eDNA extracted from influent wastewater and activated sludge supernatant samples yielded higher concentrations (12.5 ± 1.9 ng mL^−1^ and 12.3 ± 1 ng mL^−1^, respectively) than after purification (9.0 ± 0.7 ng mL^−1^ and 5.6 ± 0.46 ng mL^−1^, respectively. An average mass loss of 42.1% through the commercial purification kit was measured. In the method, only one commercial extracting kit was used and assessed for eDNA loss during purification. For future works, other commercial kits or classical purification methods might be assessed for the isolation of free-floating eDNA from low-content water samples.

IIt is paramount to note that the eDNA yields were substantial for the three water matrices. These three-floating eDNA extracts fulfilled the minimal requirements (mass ≥ 1 μg, concentration ≥ 20 ng μL^−1^ volume ≥ 20 μL) in terms of quantity and quality requested for qPCR and high-throughput metagenomics analysis. Much higher yields than other isolation methods used in this and other studies were obtained: from *ca.* 3-fold more than with classical CTAB and PCI methods up to as high as *ca.* 2000-fold higher than direct precipitation with isopropanol (**Table 3**). This method together with the one of ^50^ are the only ones showing eDNA yield and recovery from wastewater samples to be used for further molecular studies.

**Table 3.** Concentrations and yields of extracellular DNA extracts obtained during this study in comparison to other methods. Legend: cetyl trimethyl ammonium bromide (CTAB), phenol:chloroform:isoamyl alcohol and ethanol (PCI)

| Method | Concentration<br>(ng mL<sup>-1</sup>) | Initial sample volume<br>(mL) | Yield<br>(ng) |
| --- | --- | --- | --- |
| *This study* | 12.3 | 1'000 | 12'323 |
| *CTAB, PCI* | 4.53 | 800 | 3'630 |
| <sup>50</sup> | 78.0 | 5 | 234 |
| *Vivaspin columns 100 kDa* | 0.27 | 500 | 136 |
| *Isopropanol precipitation* | 0.13 | 50 | 6.5 |
| *DNA extraction kit* | ND <sup>a</sup> | 50 | ND |

<sup>a</sup> ND means notdetected

Different studies aim at isolating eDNA from different types of environmental samples and matrices. Water qualities and matrices (like biofilms, sediments, effluent water, influent wastewater, freshwater, permafrost, cattle rumen, *etc.*) and biogeographical areas do play an important role in the quantity of eDNA that can be retrieved. Thus, complicating the comparison between already published methods for isolating eDNA.

### Mild enzymatic post-treatment with proteinase K is necessary to release eDNA from bound extracellular polymeric proteins found in activated sludge

The free-floating eDNA extracted from activated sludge was characterized by an intense band at the top of the agarose gel. This suggested that eDNA was entrapped in a protein mesh that did not allow it to migrate through the gel **(Figure 4a)**. After mild enzymatic treatment of these residual proteins bound to the eDNAs with proteinase K **(Figure 4b)**, it was observed that the purified eDNA templates were able to run through the gel. These were characterized by a distribution of fragments sizes that ranged from 0.5 kbp to >20 kbp. For comparison, the iDNA that was extracted in parallel using commercial kits exhibited DNA fragments that also ranged from less than 0.5 kbp to >20 kbp. Both DNA fractions dsplayed similar fragment size distribution.

**Figure 4.**
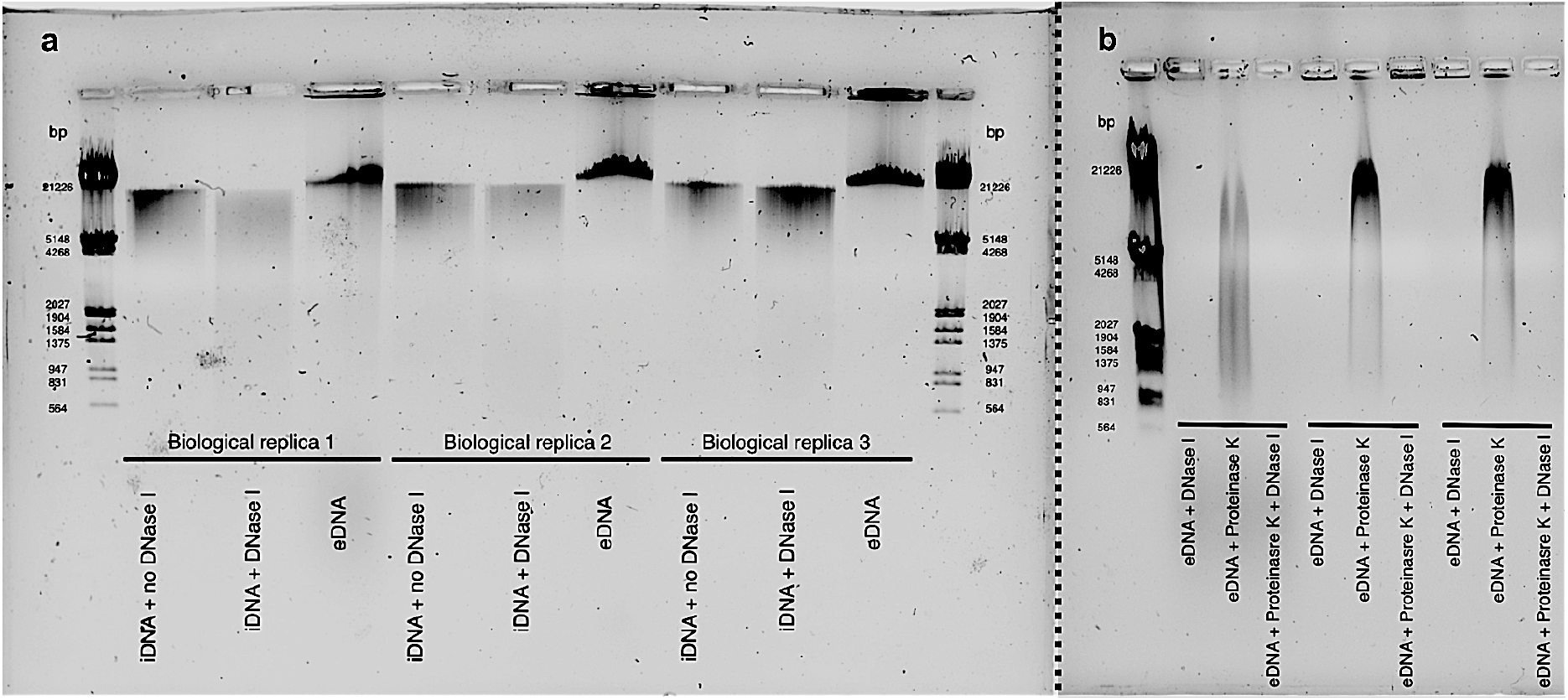
The impact of enzymatic post-treatment of raw eDNA extracts with proteinase K to release the eDNA from extracellular protein mesh: **(a)** Agarose gel from three different biological replicates showing the fragment size distributions of intracellular DNA (iDNA) obtained without any preliminary treatment, of iDNA obtained after enzymatic pre-treatment of cells with DNase I prior to cell lysis and DNA extraction, and ofthe raw extract of free-floating extracellular DNA (eDNA) obtained from from centrifuged and filtered activated sludge supernatant samples. **(b)** Agarose gel of free-floating eDNA after either pre-treatment of cells with DNase I, post-treatment of eDNA extract with proteinase K, or a combination of both pre-and post-treatments.

Assuming that the average size of genes present in bacterial genomes is 1.5 kb long, eDNA was suggested to be large enough to contain multiple ARGs and MGEs ^51,52^. No significant differences on band intensity could be observed between untreated biomass and treated with DNase I prior to intracellular DNA extraction (**Figure 4b**). This suggests a low eDNA content bound to biomass when DNA is extracted from 0.25 g of activated sludge. The extracted pools of free-floating eDNA fragments were suitable for further molecular analyses since purity (1.76 ± 0.02) was close to optimal the optimum absorbance ratio A_260 nm_ / A_280 nm_ of 1.8.

The electrophoretic behavior of the non-treated eDNA extract has been observed when eDNA has been extracted from biofilms^53,54^. Raw eDNA extracts have been hampered in their PCR amplifications. This suggests that eDNA may be co-extracted in a mesh of extracellular polymeric substances (EPS) that must be degraded in order to release the eDNA template prior tomolecular analyses. Further examination of the chemical composition of raw eDNA extracts are necessary to fully understand their characteristics and peculiarities. Mild enzymatic treatment with proteinase K is used in many commercial kits during DNA extraction in order to remove residual proteins from DNA extracts. Its use is also recommended to eliminate nucleases that degrade nucleic acids. Even if some studies have suggested that the use of high proteinase K concentrations (was not the case here) could fragment DNA ^55,56^, an heterogeneous eDNA fraction that would represent the whole cell-free aqueous solution was expected.

### Cell lysis is not induced during the isolation of free-floating eDNA

Cell lysis measurements and control were conducted on the processing of activated sludge since exhibiting the highest microbial density among the wastewater environment samples. Thus, activated sludge was considered as more prone to potential cell lysis and cross-contamination of the free-floating eDNA extract with iDNA residues. Flow cytometry was used to measure the total cells counts after each biomass processing step in the developed protocol (**Figure 5a**) together with the relative abundances of live/dead cells measured after stainings with SYBR-Green and propidium iodide fluorophores, respectively (**Figure 5b**). Centrifugation did not impact the viability of the cells, thus no cells lysis. After filtration of centrifuged activated sludge supernatant through the 0.45 μm membrane filter, the number of flow cytometry events mL^−1^ was below 10; after second filtration on 0.2 μm, below 1 event. The diluted activated sludge control that consisted of non-centrifuged nor non-filtered activated sludge sample had an average of 1606 events mL^−1^. Cell viability on cells retained in the membrane filters was assumed not to be impaired and maintain its robustness ^14,57^.

**Figure 5.**
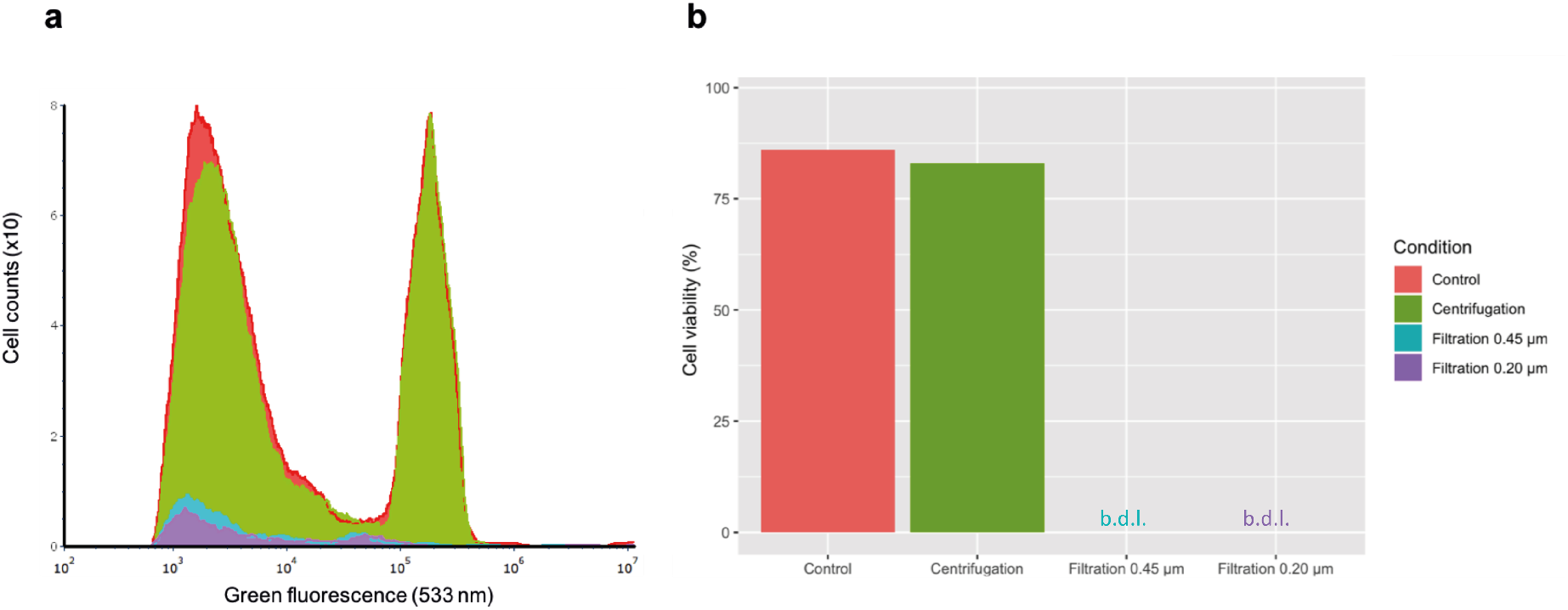
**(a)** Direct comparison of the green fluorescence histograms of activated sludge supernatant water samples with the different protocol extraction steps. Cell counts corresponds to number of events in 50 μL **(b)** Live/cell staining showing cell viability of samples after the different protocol extraction steps. **bdl:** below detection limit: cell lower limit was achieved when both filtrations were applied.

Initial cell viability was preserved and bacterial cells were removed upfront by centrifugation and filtration (depending on the type of biomasses targeted, even smaller filter pore sizes than 0.2 μm could be used). Besides high yield, the free-floating eDNA fraction was therefore of high quality, being deprived from iDNA contamination.

Overall, the method did not induce cell lysis during the extraction, meaning that a cell-free and iDNA-free free-floating eDNA fraction was isolated. This quality control ensured that in the eDNA fraction there was no genomic DNA contamination caused by the extraction. We demonstrated for the first time the isolation of free-floating eDNA at high yield and high quality from complex microbial community matrices of activated sludge, in across a workflow exempt of cell lysis.

### Free-floating eDNA displays lesser ARG copies than iDNA, but some genes exhibit similar copy numbers in both DNA fractions

qPCR results highlighted that molecular analysis of five selected ARGs (*sul1*, *sul2*, *ermB*, *qnrS*, *bla_CTXM_*) and one MGE (*intl1*) could efficiently be performed from both the iDNA and free-floating eDNA fractions of wastewater environments, being effluent wastewater in this case (**Figure 6**). All genes tested but *bla_CTXM_* were detected in both DNA fractions. The free-floating eDNA template harbored a lower number of ARG and MGE copies than iDNA, with a significant difference (Δ) of 0.87 ± 0.32 log_10_ gene copies (p<0.005) across the gene panel.

**Figure 6.**
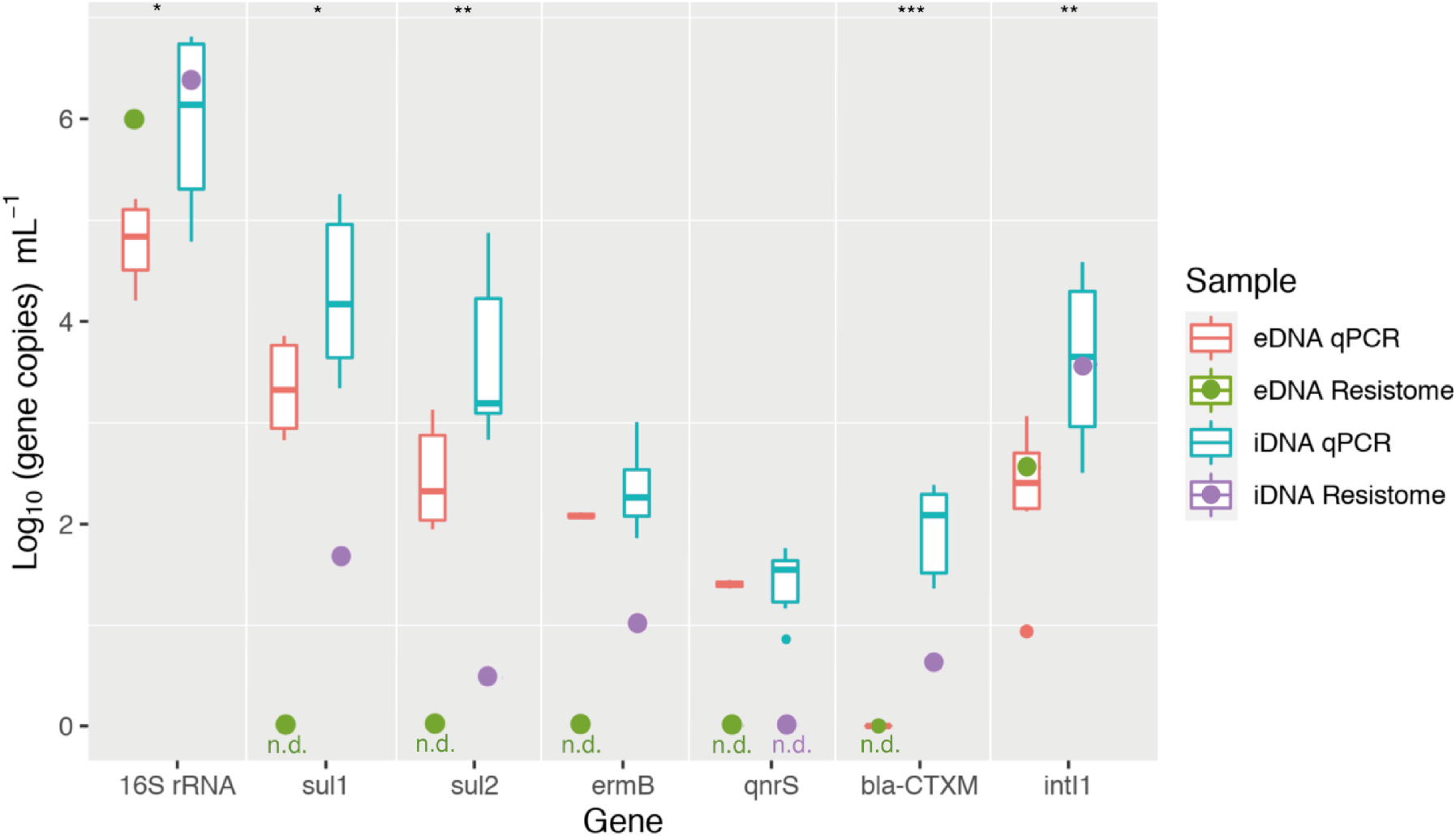
Quantitative PCR results from effluent water and number of primary aligned reads from the resistome analysis from activate sludge samples assessing the differences between DNA fractions (eDNA and iDNA). Values are displayed as Log_10_ gene copies mL-1 from a selected panel of *16S rRNA* gene, five antibiotic resistance genes (*sul1, sul2, ermB, qnrS and bla_CTXM_*), and one mobile genetic element (*intI1*). a p<0.05(*), p<0.005 (**), p<0.0005 (***). Resistome 0 values meant non detected primary aligned reads (n.d.).

At individual gene level, the free-floating eDNA displayed a significantly lower log-based gene copy number of both the bacterial *16S rRNA* gene (Δ = 1.2 ± 0.43 log_10_ gene copies) and the ARGs (difference values given hereafter) *vs.* iDNA; with the exception of the macrolides (Δ *ermB* = 0.27 ± 0.24 log_10_ gene copies) and fluoroquinolone (Δ *qnrS* <0.1 log_10_ gene copies) resistance genes that displayed similar copy numbers in iDNA and eDNA. Conversely, sulfonamides (Δ *sul1* = 0.94 ± 0.33; Δ *sul2* = 1.11 ± 0.32 log_10_ gene copies mL^−1^) and β-lactamase (Δ *bla_CTXM_* = 1.92 ± 0.25 log_10_ gene copies mL^−1^) resistance genes and the integrase type I (Δ *intI1* = 1.32 ± 0.38 log_10_ gene copies mL^−1^) MGE were significantly lesser in the free-floating eDNA (*i.e.*, outside the cell) than iDNA (*i.e.*, inside the cell).

Macrolide resistance genes (*ermB*) have been described to be embedded in transposon-like elements such as Tn551, Tn552, Tn4001 or Tn4003 conferring resistance to macrolide-lincosamide-steptogramin B (MLS) ^58^. Fluoroquinolones resistance genes have been reported to be plasmid-borne since 1998^59^. Most of *qnr* genes have been detected in *Entereobacteriaceae*, being *qnrS* gene prevalent both in environmental strains ^60^ and in non-conjugative plasmids harbouring *mob* genes allowing their mobilization^61^. Those ARGs happen to commonly be found in MGEs, principal subsystem component of the eDNA fraction (**Figure 8**).

The genes encoding resistance to sulfonamides (*sul1* and *sul2*) were the most abundant ARGs in both iDNA and eDNA fractions from the selected panel, which happens to be one of the most used antibiotics for systemic use in hospitals in The Netherlands (National Institute for Public Health and the Environment, 2019). Such high level is in agreement to other studies that have measured ARGs from total DNA extracted from activated sludge biomasses and effluent water ^63,64^. Lower number of ARGs copies have already been observed in the eDNA fraction when compared to the iDNA from sludge using a protocol for extracting eDNA from marine sediments ^51,65^.

We made a key breakthrough with our study by making the direct molecular comparison of free-floating eDNA and iDNA fractions possible, such as on the very important problematic of ARGs and MGEs in the contexts of environmental and public health protection.

Lower values measured in the eDNA fraction can arise from different possible causes. It may be considered that DNA fragments of the genes are less released outside cells, thus interrogating whether *sul1/2* and *bla_CTXM_* genes are less transferred on plasmids thus less mobile. It can also suggest that iDNA, when released because of cell decay or active release mechanisms, may be degraded by nucleases present in both DNA fractions. Another possibility is the degradation of DNA fragments that carried these genes by microorganisms, but DNA degradation by microbes is likely not specific at gene level. All in all, the reason for this difference between compositions of iDNA and free-floating eDNA fractions need to get investigated. Our isolation method provides the analytical key to investigate it.

What it is highly remarkable is that ARGs can still be detected in significant amounts (100-1000 gene copies mL^−1^) in the eDNA fraction that is fully exposed to the environment. A plausible explanation why eDNA persists in aqueous environments is that it is clothed with a mesh of extracellular proteins (shown here by the proteinase K post-treatments). Other causes of protection against degradation by nucleases can relate to eDNA being locked within organic and inorganic aggregates, adsorbed onto mineral matrices, or integrated in viral genomes ^1^.

The number of primary reads from the simplicate resistome analysis by metagenomics of free-floating eDNA retrieved from activated sludge (see next section in details) were also displayed on **figure 6** for comparison to qPCR data. Genes selection was done based on the panel of genes selected for qPCR. Since we cannot compare both water samples (resistome was done on activated sludge samples and qPCR on effluent water samples), this was done to verify if there was a tendency in terms of quantities between DNA fractions followed by both ARGs and MGEs detection methods. The resistome dataset showed a lower log_10_ number of ARG reads when compared to qPCR data. This is due to lower sensibility of metagenomics when compared to PCR-based methods on wastewater samples ^66^. Collectively our and previous studies highlight the need to combine molecular methods for an accurate analysis of ARGs and MGEs in wastewater. It is not all about high-throughput sequencing. Metagenomics provides high resolution on the diversity of genes and there alleles. qPCR provide high sensitivity for the detection of ARGs and MGEs. Again, our study provides a strong new insight on the separation and comparison of free-floating eDNA fractions from complex environmental biological samples.

Genes displayed in **figure 6** (*sul1, sul2, ermB, qnrS* and *bla_CTXM_*) corresponded to less than 2% of the total primary ARGs reads found in the iDNA resistome. From the activated sludge resistome, only the number of reads of the universal *16S rRNA* and ubiquitous *intI1* genes were close to the ones measured by qPCR in both DNA fractions. When the trend in gene copies per mL was compared, iDNA fractions contained more number of primary aligned reads from the selected panel of genes than the eDNA fraction. This corresponds on what was observed for qPCR results.

The resistome analysis fully depends on the databases selected for gene mapping. ARG sequences that are highly dissimilar to reference sequences deposited in databases cannot be detected, thus not resulting in hits and not appearing in final results ^67^.

### Metagenomics allowed for a broad and high-resolution analysis and comparison of the resistome, mobilome and microbiome compositions of the free-floating eDNA and microbial iDNA fractions

The quantity and quality of the eDNA and iDNA templates separately retrieved from the complex activated sludge enabled a high-throughput metagenomics analysis with high resolution on the mobilome, resistome and microbiome compositions (**Figure 7**). The number of reads used for metagenomic analysis are summarized in **Table 4**. The absolute read counts of specific genes were then translated into relative abundances when normalized by the total number of primary aligned reads after filtration per database selected.

**Table 4.** Number of reads obtained at every step of the dry-lab workflow used to analyse and compare the metagenomes of free-floating extracellular DNA (eDNA) and intracellular DNA (iDNA) fractions isolated from activated sludge.

| Bioinformatic step | Free-floating eDNA | iDNA |
| --- | --- | --- |
| Number of raw reads out of sequencer | 14'127'626 | 14'788'250 |
| Number of reads after quality control | 7'830'000 | 6'480'000 |
| Number of MEGAREs<sup>a</sup> primary aligned reads before samclip<sup>b</sup> | 188 | 9'725 |
| Number of MEGAREs<sup>a</sup> primary aligned reads after samclip<sup>b</sup> | 82 | 1'571 |
| Number of ACLAME<sup>c</sup> primary aligned reads before samclip<sup>b</sup> | 546'007 | 1'183'308 |
| Number of ACLAME<sup>c</sup> primary aligned reads after samclip<sup>b</sup> | 180'860 | 315'225 |

<sup>a</sup> Antimicrobial Database for High-Throughput Sequencing.
<sup>b</sup> Tool designed to remove soft and hard clipped alignments.
<sup>c</sup> Database dedicated to the collection and classification of mobile genetic elements (MGEs).

**Figure 7.**
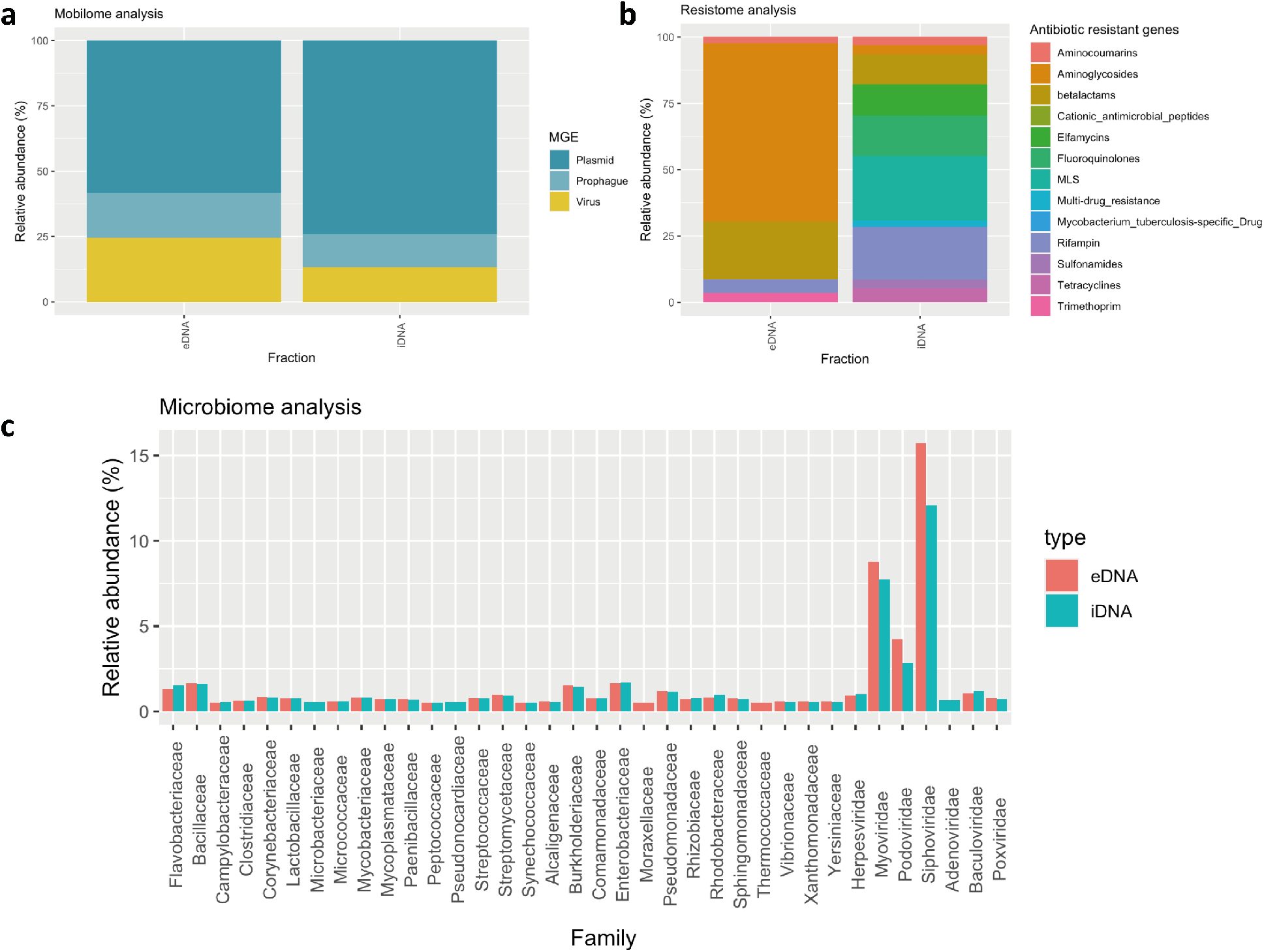
Metagenome analysis from eDNA and iDNA fractions obtained from an activated sludge sample **(a)** Mobilome relative abundance. **(b)** Resistome relative abundance. **(c)** Microbiome relative abundance showing bacterial and viral families, whose population composition was >0.5%.

*Mobilome* results showed that both the free-floating eDNA and the iDNA fractions contained a higher relative abundance of plasmids (58% and 74%, respectively) when compared to viruses and prophages (41% and 26%, respectively) (**Figure 7a**). In the eDNA fraction, the relative abundance of plasmids was significantly lower than the viral fraction. This might be explained by extracellular plasmids exposure to and degradation by environmental nucleases. The presence of microorganisms may also enhance the degradation rate of extracellular DNA plasmids ^68^; Thus, increasing the relative abundance of viral units in the extracellular DNA fraction.

Detailed mobilome compositions are summarized in the supplementary material **Figure S2** for both free-floating eDNA and iDNA. From the eDNA fraction, integrons, integrative conjugative elements (ICE) and insertion sequences (IS) were analyzed being *intI1* (92%) **(Figure S2b)**, actinomycete conjugative integrative element (AICE) (35%) and SXT/R391 (30%) **(Figure S2c)** and IS200 (38%) **(Figure S2d)** the most abundant components in their category, respectively.

Acquisition of antibiotic resistances does not only depend on the uptake or exchange of ARGs themselves but also on the uptake of MGEs such as IS, transposons, and gene cassettes/integrons that carry and facilitate the uptake of ARGs into cells ^58^. IS and integrons, especially the class I, can influence the level of transcription or indirectly affect genes involved in their regulation or in the modulation of resistance levels ^69,70^. Class 1 integrase *intI1* has been considered as a good proxy for anthropogenic pharmaceutical pollution being usually linked to genes conferring resistance to antibiotics^35^. A fraction of 80% of commercial antibiotics are produced with bacteria from the genus *Streptomyces* and rare Actinomycetes such as *Actinomadura* ^71^. It has been observed that ARGs are associated with AICEs, being able to have a role in horizontal gene transfer at least between actynomycetes ^72^. SXT/R391 is a type of ICE closely related to *Enterobacteriaceae*, one of the predominant bacterial families present in our samples (**Figure 7c)**. Species of this family have been described to majorly contain four types of ARGs (*straA/B, sul2, floR* and *dfrA*) ^73^. IS200 elements contain a single gene and are found in bacterial and archaeal species, but it has been described as a rarely transposable element^74^. MGEs have an effect on the ARGs mobility as well as on the regulation at the expression level of these genes^75^.

*Resistome* results showed more distinct patterns among DNA fractions (**Figure 7b**). We are, for the first time, showing the profile of the extracellular DNA resistome from an activated sludge sample. The free-floating eDNA fraction harbored a total of 12 different ARGs from 5 different antibiotic resistance families. There was a clear relative dominance of aminoglycosides (67%) and β-lactams (22%) resistant genes on the eDNA.

A total of 289 different ARGs from 15 different antibiotic resistance families were found in the iDNA fraction. This matches with the latest work of Jia et al. (2020) who detected 297 different genes from 17 different families in water and water filters from a drinking water treatment plant. MLS dominated (24%), followed by rifampin (20%), fluoroquinolones (15.1%) and elfamycins (11.7%) among other resistance genes on the iDNA.

Most of the ARGs are enclosed inside bacterial cells while some specific types may endure floating or are bound to matrices in activated sludge samples, even if in low quantities. Low amount of free floating ARGs suggests that natural transformation may not be the main mechanisms through which gene transfer and exchange does occur ^51^. However, this smaller amount of extracellular ARGs does not preclude any possibility to be taken up by naturally competent bacteria in complex cultures via transformation processes.

*Microbiome* profiles were more conserved **(Figure 7c)**. Operational taxonomic units (OTUs) found in the extracellular fraction are genomic fragments that at some point have been released from lysed or necropsied cells. These genes are then easier to be degraded than plasmid or viral DNA, due to exposed fragmented double-stranded DNA to nucleases. Thus, it is more complicated to observe population of microorganisms migrations during the wastewater treatment process. Moreover, some studies have suggested that DNA adsorbed to soil or EPS matrices are protected from degradation by environmental nucleases, indicating that genes on plasmid may be better preserved than genomic DNA^77,78^. In this case, it can be observed that only some specific bacterial and viral families are exclusive from one or another DNA fraction if we consider the 0.5% relative abundance cutoff. Examples are *Microbacteriacae, Pseudonocardiacae* and *Adenoviridae* for the iDNA fraction and *Moraxellaceae* and *Thermococcaceae* for the eDNA **(Figure 7c)**. The *Moraxellaceae* family is known to be an inductor of activated sludge flocculation ^79^, which made no sense that was only present in the extracellular DNA fraction. When the raw data was checked, it was observed that its relative abundance in the intracellular DNA was 0.48%. This is the reason why it did not appear in the graph (>0.5% cutoff). The same situation was observed with the other bacterial and viral families that were found only in one of the DNA fraction. Thus, this meant that no significant population differences could be observed between families and DNA fractions from activated sludge samples. We hypothetize that some eDNA may come directly from influent water but mainly from activated sludge. This eDNA from microbial populations could have been directly obtained from decaying populations in the activated sludge tank making this method an efficient way to track decay phenomena in activated sludge at population resolution level.

### Genomic functional subsystems characterization highlighted different patterns between free-floating eDNA and iDNA

Results of assigning the metagenomics sequences to functional categories (level 1 of subsystem) are shown in **Figure 8**. Approximately 40% and 50% of the total predicted proteins with known functions (808’680 and 6’771’570, respectively) from the eDNA and iDNA metagenomes showed matches against the subsystems database on the MetaGenome Rapid Annotation with Subsystem Technology (MG-RAST) server^45^.

**Figure 8.**
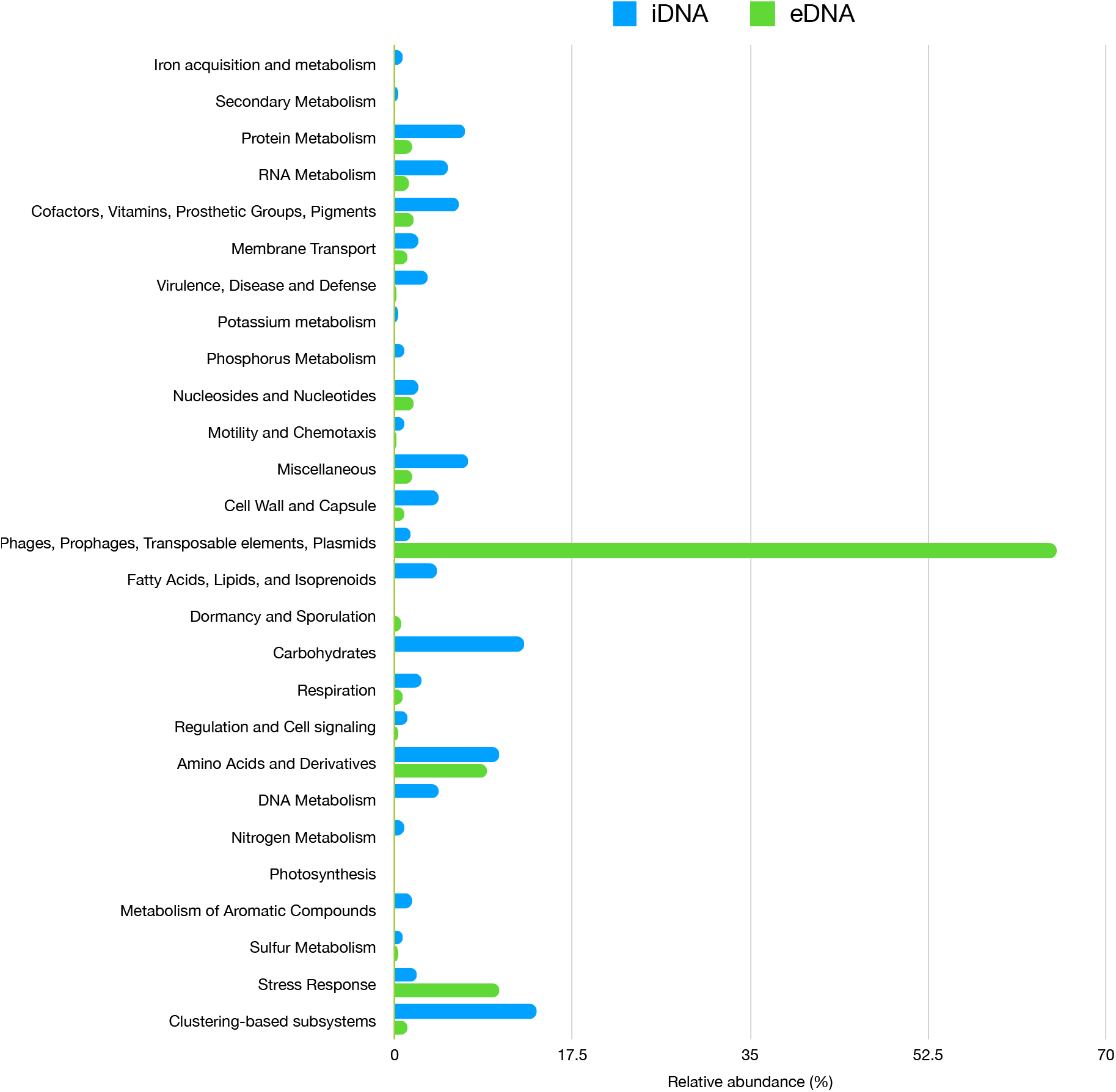
Subsystem classification of the functional genes identified from the metagenome of the free-floating extracellular DNA fraction (eDNA, *green*) fraction and the intracellular DNA (iDNA, *blue*) fractionusing MG-RAST for mapping and computation of relative abundances (%)

Interestingly, patterns in the data confirmed that eDNA metagenome was accomplished predominantly by phages, prophages, transposable elements and plasmids (65.1%, 526’534 hits). The same category corresponded to a residual 1.6%, 108’713 hits on the iDNA metagenome. iDNA fraction showed an heterogeneous distribution, classified mainly by clustering-based subsystems (13.94%, 944’394 hits) carbohydrates (12.8%, 865’220 hits), and amino acids and derivatives (9.13%, 701’175 hits) sublevels, among others. Clustering-based sybsystems contain such functions as proteosomes, ribosomes and recombination-related clusters ^80^, which are localized in microorganisms cytoplasm (iDNA). Overall, the free-floating eDNA is confirmed to be a pool of DNA mainly composed by MGEs. These do not necessarily harbor ARGs in their structural components. Thus, making eDNA a potential DNA source for horizontal gene transfer mechanisms. This could generate new microbial diversity as well as mobilize ARGs^23^.

## Conclusion

The implementation of DEAE chromatography promoted an efficient isolation of free-floating eDNA at high yield and quality from complex matrices of WWTP environments from influent wastewater to activated sludge and effluent water samples. It provides a rightful alternative to classical methods to extract eDNA for further molecular quantification at high sensitivity (with qPCR) and profiling at high throughput and resolution (with metagenome sequencing) of ARGs and MGEs. Thus, allowing both the study from a metagenomics point of view to describe the diversity of xenogenic elements and from a quantitative PCR approach to analyze representative panels of selected genes. Quantitative results showed from 1.8 to 3.8 log_10_ number of ARGs copies in the free-floating eDNA fraction. It showed significant differences between eDNA and iDNA fractions, especially for those most abundant ARGs such as sulfonamides (0.93 and 1.11 log_10_ difference for *sul1* and *sul2*, respectively). The metagenomics analysis of the microbiome compositions resulted in non-significant variation in taxonomic families between DNA fractions, suggesting that a big fraction of free DNA found in activated sludge may derive from the flocs. The mobilome and resistome primary aligned reads from the free-floating eDNA harbored lower numbers than iDNA but could still from a potential source of ARGs and MGEs for natural transformation. Interestingly, subsystems classifications showed that eDNA fraction is mainly composed by MGEs (65.1%), confirming the hypothesis that eDNA is a pool of MGEs for horizontal gene transfer. Further studies on the conditions promoting horizontal gene transfer in complex biological systems will be needed in order to elucidate the actual natural transformation rate. Molecular quantification and metagenomics profiling from free-floating eDNA after isolation using our novel, chromatographic method will be powerful to support the design of new risk assessments on xenogenic elements like ARG, MGE, but also residual GMO materials across urban and natural water systems. It will support the delineation of measures by water authorities to adopt at WWTPs to remove them, and safeguard environmental and public health.

## Conflict of interest statement

The authors declare no conflict of interest.

## Authors’ contributions

DCF designed the study with inputs of MvL, TA and DGW. DCF performed the experimental investigations. DCF wrote the manuscript with direct contribution, edits, and critical feedback by all authors.

## Acknowledgements

We are very grateful to Pilar de la Torre Cortés from the Industrial Microbiology Section at TU Delft for preparing the libraries and sequencing our DNA samples. This work is part of the research project “Transmission of Antimicrobial Resistance Genes and Engineered DNA from Transgenic Biosystems in Nature” (Targetbio) funded by the programme Biotechnology & Safety (grant no. 15812) of the Applied and Engineering Sciences Division of the Dutch Research Council (NWO).

## Supplementary information

**Table S1.** 16S rRNA, ARGs and MGE synthetic DNA fragments used from ResFinder for generating standard curves used for qPCR analysis

**Table S2.** Primers used in this study.

**Table S3.** Mobilome number of primary reads after INTEGRALL, ISfinder and ICEberg alignment

**Figure S1.** Control chromatogram (λ=260 nm) with **(a)** solution containing pure plasmid pHT01 (0.73 ng μL^−1^, 10 mL) **(b)** Raw eDNA from filtered activated sludge water sample (900 mL) chromatogram (λ=260 nm).

**Figure S2.** Mobilome analysis from eDNA and iDNA fractions obtained from an activated sludge sample **(a)** Integrative, conjugative (ICE), mobilizable (IME) and cis-Mobilizable elements (CIME) relative abundance. **(b)** Integrons relative abundance. **(c)** Global mobile genetic elements relative abundance. **(d)** Integrative conjugative elements (ICE) relative abundance and **(e)** insertion sequences relative abundance.

## Notes

### Competing Interest Statement

The authors have declared no competing interest.

## References

1. Torti, A., Lever, M. A. & Jørgensen, B. B. Origin, dynamics, and implications of extracellular DNA pools in marine sediments. Mar. Genomics 24, 185–196(2015).

2. Mao, D. et al. Persistence of extracellular DNA in river sediment facilitates antibiotic resistance gene propagation. Environ. Sci. Technol. 48, 71–78 (2014).

3. Vuillemin, A. et al. Preservation and significance of extracellular DNA in ferruginous sediments from Lake Towuti, Indonesia. Front. Microbiol. 8, 1–15 (2017).

4. Lorenz, M. G. & Wackernagel, W. Bacterial gene transfer by natural genetic transformation in the environment. Microbiological reviews 58, (1994).

5. Alekseeva, L. A. et al. Alteration of the exDNA profile in blood serum of LLC-bearing mice under the decrease of tumour invasion potential by bovine pancreatic DNase I treatment. PLoS One 12, 1–21 (2017).

6. Thomsen, P. F. & Willerslev, E. Environmental DNA - An emerging tool in conservation for monitoring past and present biodiversity. Biol. Conserv. 183, 4–18 (2015).

7. Shogren, A. J. et al. Modelling the transport of environmental DNA through a porous substrate using continuous flow-through column experiments. J. R. Soc. Interface 13, (2016).

8. Majaneva, M. et al. Environmental DNA filtration techniques affect recovered biodiversity. Sci. Rep. 8, 1–11 (2018).

9. Dominiak, D. M., Nielsen, J. L. & Nielsen, P. H. Extracellular DNA is abundant and important for microcolony strength in mixed microbial biofilms. Environ. Microbiol. 13, 710–721 (2011).

10. Rusanowska, P., Cydzik-Kwiatkowska, A. & Wojnowska-Baryła, I. Microbial Origin of Excreted DNA in Particular Fractions of Extracellular Polymers (EPS) in Aerobic Granules. Water. Air. Soil Pollut. 230, (2019).

11. Weissbrodt, D. G., Neu, T. R., Kuhlicke, U., Rappaz, Y. & Holliger, C. Assessment of bacterial and structural dynamics in aerobic granular biofilms. Front. Microbiol. 4, 1–18 (2013).

12. Tetz, V. V. & Tetz, G. V. Effect of extracellular DNA destruction by DNase I on characteristics of forming biofilms. DNA Cell Biol. 29, 399–405 (2010).

13. Tang, L., Schramm, A., Neu, T. R., Revsbech, N. P. & Meyer, R. L. Extracellular DNA in adhesion and biofilm formation of four environmental isolates: A quantitative study. FEMS Microbiol. Ecol. 86, 394–403 (2013).

14. Wu, J. & Xi, C. Enzymatic method for extracting extracellular DNA in biofilm matrix. Cold Spring Harb. Protoc. 5, (2010).

15. Merod, R. T. & Wuertz, S. Extracellular polymeric substance architecture influences natural genetic transformation of Acinetobacter baylyi in biofilms. Appl. Environ. Microbiol. 80, 7752–7757 (2014).

16. Flemming, H. C. & Wingender, J. The crucial role of extracellular polymeric substances in biofilms. Nat. Rev. Microbiol. 8, 623–633 (2010).

17. Nagler, M., Insam, H., Pietramellara, G. & Ascher-Jenull, J. Extracellular DNA in natural environments: features, relevance and applications. Appl. Microbiol. Biotechnol. 102, 6343–6356 (2018).

18. Okshevsky, M. & Meyer, R. L. Evaluation of fluorescent stains for visualizing extracellular DNA in biofilms. J. Microbiol. Methods 105, 102–104 (2014).

19. Cheng, M., Cook, A. E., Fukushima, T. & Bond, P. L. Evidence of compositional differences between the extracellular and intracellular DNA of a granular sludge biofilm. Lett. Appl. Microbiol. 53, 1–7 (2011).

20. Stoeckle, M. Y., Soboleva, L. & Charlop-Powers, Z. Aquatic environmental DNA detects seasonal fish abundance and habitat preference in an urban estuary. PLoS One 12, 1–15 (2017).

21. Corinaldesi, C., Tangherlini, M., Manea, E. & Dell’Anno, A. Extracellular DNA as a genetic recorder of microbial diversity in benthic deep-sea ecosystems. Sci. Rep. 8, 1–9 (2018).

22. Claverys, J. P., Martin, B. & Polard, P. The genetic transformation machinery: Composition, localization, and mechanism. FEMS Microbiol. Rev. 33, 643–656 (2009).

23. Jørgensen, T. S., Kiil, A. S., Hansen, M. A., Sørensen, S. J. & Hansen, L. H. Current strategies for mobilome research. Front. Microbiol. 5, 1–6 (2014).

24. Calderón-Franco, D., Lin, Q., Loosdrecht, M. C. M. Van Abbas, B. & Weissbrodt, D. G. Anticipating Xenogenic Pollution at the Source : Impact of Sterilizations on DNA Release From Microbial Cultures. Front. Bioeng. Biotechnol. 8, 1–13 (2020).

25. Wang, D. N. et al. A new adsorption-elution technique for the concentration of aquatic extracellular antibiotic resistance genes from large volumes of water. Water Res. 92, 188–198 (2016).

26. Nagler, M., Podmirseg, S. M., Griffith, G. W., Insam, H. & Ascher-Jenull, J. The use of extracellular DNA as a proxy for specific microbial activity. Appl. Microbiol. Biotechnol. 102, 2885–2898 (2018).

27. Katevatis, C., Fan, A. & Klapperich, C. M. Low concentration DNA extraction and recovery using a silica solid phase. PLoS One 12, 1–14 (2017).

28. Krajacic, M., Ravnikar, M., Štrancar, A. & Gutiérrez-Aguirre, I. Application of monolithic chromatographic supports in virus research. Electrophoresis 38, 2827–2836 (2017).

29. Moore, D. & Dowhan, D. Purification and Concentration of DNA. Curr. Protoc. Mol. Biol.(2002).

30. Prest, E. I., Hammes, F., Kötzsch, S., van Loosdrecht, M. C. M. & Vrouwenvelder, J. S. Monitoring microbiological changes in drinking water systems using a fast and reproducible flow cytometric method. Water Res. 47, 7131–7142 (2013).

31. Hammes, F. et al. Flow-cytometric total bacterial cell counts as a descriptive microbiological parameter for drinking water treatment processes. Water Res. 42, 269–277 (2008).

32. Pinel, I. S. M., D.H., M., Vrouwenvelder, J. S. & Van Loosdrecht, M. C. M. Bacterial community dynamics and disinfection impact in cooling water systems. Water Res. 172, 115505 (2020).

33. Boulos, L., Prévost, M., Barbeau, B., Coallier, J. & Desjardins, R. LIVE/DEAD(®) BacLight(TM): Application of a new rapid staining method for direct enumeration of viable and total bacteria in drinking water. J. Microbiol. Methods 37, 77–86 (1999).

34. Pallares-Vega, R. et al. Determinants of presence and removal of antibiotic resistance genes during WWTP treatment: A cross-sectional study. Water Res. 161, 319–328 (2019).

35. Ma, L., Li, A. D., Yin, X. Le & Zhang, T. The Prevalence of Integrons as the Carrier of Antibiotic Resistance Genes in Natural and Man-Made Environments. Environ. Sci. Technol. 51, 5721–5728 (2017).

36. Andrews, S. FastQC: a quality control tool for high throughput sequence data. (2010).

37. Bolger, A. M., Lohse, M. & Usadel, B. Trimmomatic: A flexible trimmer for Illumina sequence data. Bioinformatics 30, 2114–2120 (2014).

38. Li, H. Aligning sequence reads, clone sequences and assembly contigs with BWA-MEM. 00, 1–3 (2013).

39. Kim, D., Song, L., Breitwieser, F. P. & Salzberg, S. L. Centrifuge: rapid and accurate classifica3on of metagenomic sequences. bioRxiv 26, 054965 (2016).

40. Leplae, R., Lima-Mendez, G. & Toussaint, A. ACLAME: A CLAssification of mobile genetic elements, update 2010. Nucleic Acids Res. 38, 57–61 (2009).

41. Liu, M. et al. ICEberg 2.0: An updated database of bacterial integrative and conjugative elements. Nucleic Acids Res. 47, D660–D665 (2019).

42. Siguier, P., Perochon, J., Lestrade, L., Mahillon, J. & Chandler*, and M. ISfinder: the reference centre for bacterial insertion sequences. Nucleic Acids Res. 34, D32–D36 (2006).

43. Moura, A. et al. INTEGRALL: A database and search engine for integrons, integrases and gene cassettes. Bioinformatics 25, 1096–1098 (2009).

44. Li, H. et al. The Sequence Alignment/Map format and SAMtools. Bioinformatics 25, 2078–2079 (2009).

45. Meyer, F. et al. The metagenomics RAST server - A public resource for the automatic phylogenetic and functional analysis of metagenomes. BMC Bioinformatics 9, 1–8 (2008).

46. R Foundation for Statistical Computing. R: a Language and Environment for Statistical Computing. http://www.R-project.org/ 2, (2018).

47. Saeki, K., Ihyo, Y., Sakai, M. & Kunito, T. Strong adsorption of DNA molecules on humic acids. Environ. Chem. Lett. 9, 505–509 (2011).

48. Feng, H. J., Hu, L. F., Mahmood, Q., Long, Y. & Shen, D. S. Study on biosorption of humic acid by activated sludge. Biochem. Eng. J. 39, 478–485 (2008).

49. Ma, H., Allen, H. E. & Yin, Y. Characterization of isolated fractions of dissolved organic matter from natural waters and a wastewater effluent. Water Res. 35, 985–996 (2001).

50. Yuan, Q. Bin et al. Redistribution of intracellular and extracellular free & adsorbed antibiotic resistance genes through a wastewater treatment plant by an enhanced extracellular DNA extraction method with magnetic beads. Environ. Int. 131, 104986 (2019).

51. Zhang, Y., Snow, D. D., Parker, D., Zhou, Z. & Li, X. Intracellular and extracellular antimicrobial resistance genes in the sludge of livestock waste management structures. Environ. Sci. Technol. 47, 10206–10213 (2013).

52. DeFlaun, M., Paul, J. & Jeffrey, W. Distribution and molecular weight of dissolved DNA in subtropical estuarine and oceanic environments. Mar. Ecol. Prog. Ser. 38, 65–73 (1987).

53. Kilic, T. et al. Biofilm characteristics and evaluation of the sanitation procedures of thermophilic Aeribacillus pallidus E334 biofilms. Biofouling 33, 352–367 (2017).

54. Chiba, A., Sugimoto, S., Sato, F., Hori, S. & Mizunoe, Y. A refined technique for extraction of extracellular matrices from bacterial biofilms and its applicability. Microb. Biotechnol. 8, 392–403 (2015).

55. Santos, A. L. F., Oliveira, C. Q. P., Arruda, G. N. P. N. & Martins, J. K. Comparison of DNA extraction using proteinase K and extraction kit: Analysis of the quality of the genetic material. J. Bras. Patol. e Med. Lab. 54, 70–75 (2018).

56. da Silva, L. E. et al. Variation in concentration of proteinase K for bovine DNA extr. Arch. Vet. Sci. 18, 15–19 (2013).

57. Tanny, G. B., Mirelman, D. & Pistole, T. Improved Filtration Technique for Concentrating and Harvesting Bacteria. Appl. Environ. Microbiol. 40, 269–273 (1980).

58. Partridge, S. R., Kwong, S. M., Firth, N. & Jensen, S. O. Mobile genetic elements associated with antimicrobial resistance. Clin. Microbiol. Rev. 31, 1–61 (2018).

59. Martínez-martínez, L., Pascual, A. & Jacoby, G. A. Quinolone resistance from a transferable plasmid. Lancet 351, 797–799 (1998).

60. Guillard, T. et al. Mobile insertion cassette elements found in small non-transmissible plasmids in Proteeae may explain qnrD mobilization. PLoS One 9, 1–8 (2014).

61. Kehrenberg, C., Hopkins, K. L., Threlfall, E. J. & Schwarz, S. Complete nucleotide sequence of a small qnrS1-carrying plasmid from Salmonella enterica subsp. enterica Typhimurium DT193. J. Antimicrob. Chemother. 60, 903 (2007).

62. ‘National Institute for Public Health and the Environment (RIVM)’. NethMap 2019. Consumption of antimicrobial agents and antimicrobial resistance among medically important bacteria in the Netherlands. (2019).

63. Rocha, J. et al. Comparison of culture-and quantitative PCR-based indicators of antibiotic resistance in wastewater, recycled water, and tap water. Int. J. Environ. Res. Public Health 16, (2019).

64. Czekalski, N., Berthold, T., Caucci, S., Egli, A. & Bürgmann, H. Increased levels of multiresistant bacteria and resistance genes after wastewater treatment and their dissemination into Lake Geneva, Switzerland. Front. Microbiol. 3, 1–18 (2012).

65. Corinaldesi, C., Danovaro, R., Anno, D. & Anno, A. D. Simultaneous Recovery of Extracellular and Intracellular DNA Suitable for Molecular Studies from Marine Sediments Simultaneous Recovery of Extracellular and Intracellular DNA Suitable for Molecular Studies from Marine Sediments. Society 71, 46–50 (2005).

66. Manaia, C. M. et al. Antibiotic resistance in wastewater treatment plants: Tackling the black box. Environ. Int. 115, 312–324 (2018).

67. Willmann, M. & Peter, S. Translational metagenomics and the human resistome: confronting the menace of the new millennium. J. Mol. Med. 95, 41–51 (2017).

68. Zhu, B. Degradation of plasmid and plant DNA in water microcosms monitored by natural transformation and real-time polymerase chain reaction (PCR). Water Res. 40, 3231–3238 (2006).

69. Jaurin, B. & Normark, S. Insertion of IS2 creates a novel ampC promoter in Escherichia coli. Cell 32, 809–816 (1983).

70. Collis, C. M. & Hall, R. M. Expression of antibiotic resistance genes in the integrated cassettes of integrons. Antimicrob. Agents Chemother. 39, 155–162 (1995).

71. Lo Grasso, L., Chillura Martino, D. & Alduino, R. Production of Antibacterial Compounds from Actinomycetes. Intech i, (2012).

72. te Poele, E. M., Bolhuis, H. & Dijkhuizen, L. Actinomycete integrative and conjugative elements. Antonie van Leeuwenhoek, Int. J. Gen. Mol. Microbiol. 94, 127–143 (2008).

73. Li, X. et al. SXT/R391 integrative and conjugative elements in Proteus species reveal abundant genetic diversity and multidrug resistance. Sci. Rep. 6, 4–12 (2016).

74. Beuzón, C. R., Chessa, D. & Casadesús, J. IS200: An old and still bacterial transposon. Int. Microbiol. 7, 3–12 (2004).

75. Depardieu, F., Podglajen, I., Leclercq, R., Collatz, E. & Courvalin, P. Modes and modulations of antibiotic resistance gene expression. Clin. Microbiol. Rev. 20, 79–114 (2007).

76. Jia, S., Bian, K., Shi, P., Ye, L. & Liu, C.-H. Metagenomic profiling of antibiotic resistance genes and their associations with bacterial community during multiple disinfection regimes in a full-scale drinking water treatment plant. Water Res. 115721 (2020). doi:10.1016/J.WATRES.2020.115721

77. Grande, R. et al. Extracellular DNA in Helicobacter pylori biofilm: A backstairs rumour. J. Appl. Microbiol. 110, 490–498 (2011).

78. Corinaldesi, C., Beolchini, F. & Dell’Anno, A. Damage and degradation rates of extracellular DNA in marine sediments: Implications for the preservation of gene sequences. Mol. Ecol. 17, 3939–3951 (2008).

79. Shchegolkova, N. M. et al. Microbial community structure of activated sludge in treatment plants with different wastewater compositions. Front. Microbiol. 7, 1–15 (2016).

80. Delmont, T. O. et al. Structure, fluctuation and magnitude of a natural grassland soil metagenome. ISME J. 6, 1677–1687 (2012).

