## Supplementary material for "A novel method to isolate free-floating extracellular DNA from wastewater for quantitation and metagenomic profiling of mobile genetic elements and antibiotic resistance genes"

Deposited as pre-print on bioRxiv

### Supplementary material

#### qPCR mix solution and reaction conditions

All ARGs and *intI1* qPCR reactions were conducted in 20 µL, including IQ$^{TM}$ SYBR green supermix BioRad 1x. Forward and reverse primers, and oligonucleotide probes (when applicable) are summarized in **Table S1 and S2.** A total of 2 µL of DNA template was added to each reaction, and the reaction volume was completed to 20 µL with DNase/RNase free Water (Sigma Aldrich, UK). All reactions (were performed in a qTOWER3 Real-time PCR machine (Westburg, DE) according to the following PCR cycles: 95ºC for 5 minutes followed by 40 cycles at 95ºC for 15 s and 60ºC for 30 s. The annealing temperature was the same for all the different reactions except for the s*ul2* and *sul1* genes. In those cases, the annealing temperatures were 61ºC and 65ºC, respectively.

In order to check the specificity of the reaction, a melting curve was performed from 65 to 95ºC at a temperature gradient of +0.5ºC $(5\ s)^{-1}$. Synthetic DNA fragments (IDT, USA) containing each of the target genes were used as a positive control to create the standard curves. Serial dilutions of gene fragments were performed in sheared salmon sperm DNA 5 µg $mL^{-1}$ (m/v) (Thermofisher, LT) diluted in Tris-EDTA (TE) buffer at pH 8.0. Every sample was analyzed in technical triplicates. Standard curves were included in each PCR plate with at least 6 serial dilutions points and in technical duplicate. An average standard curve based on a standard curve from every run was created for every gene set. Gene concentration values were then calculated from the aforementioned curve.

**Table S1.** 16S rRNA, ARGs and MGE synthetic DNA fragments ued from ResFinder to generate standard curves for qPCR

| ***Gene*** | ***Sequence*** |
|---|---|
| ***16S rRNA*** | *ACTCCTACGGGAGGCAGCAGTGGGGAATATTGCACAATGGGCGCAAGCCTGATGCAGCCATGCCGCGTGTATGAAGAAGGCCTTCGGGTTGTAAAGTACTTTCAGCGGGGAGGAAGGGAGTAAAGTTAATACCTTTGCTCATTGACGTTACCCGCAGAAGAAGCACCGGCTAACTCCGTGCCAGCAGCCGCGGTAAT* |
| ***qnrS*** | *GACGTGCTAACTTGCGTGATACGACATTCGTCAACTGCAAGTTCATTGAACAGGGTGATATCGAAGGCTGCCACTTTGATGTCGCAGATCTTCGTGATGCAAGTTTCCAACAATGCCA* |
| ***intI1*** | *GCCTTGATGTTACCCGAGAGCTTGGCACCCAGCCTGCGCGAGCAGCTGTCGCGTGCACGGGCATGGTGGCTGAAGGACCAGGCCGAGGGCCGCAGCGGCGTTGCGCTTCCCGACGCCCTTGAGCGGAAGTATCCGCGCGCCGGGCATTCCTGGCCGTGGTTCTGGGTTTTTGCGCAGCACACGCATTCGACCGATC* |
| ***sul1*** | *CGCACCGGAAACATCGCTGCACGTGCTGTCGAACCTTCAAAAGCTGAAGTCGGCGTTGGGGCTTCCGCTATTGGTCTCGGTGTCGCGGAAATCCTTCTTGGGCGCCACCGTTGGCCTTCCTGTAAAGGATCTGGGTCCAGCGAGCCTTGCGGCGGAACTTCA* |
| ***sul2*** | *TGGAGGCCGGTATCTGGCGCCAGACGCAGCCATTGCGCAGGCGCGTAAGCTGATGGCCGAGGGGGCAGATGTGATCGACCTCGGTCCGGCATCCAGCAATCCCGACGCCGCGCCTGTTTCGTCCGACACAGAAATCGCGCGTATCGCGCCGGTGCTGGACGCGCTCAAGGCAGATGGCATTCCCG* |
| ***ermB*** | *AAAACTTACCCGCCATACCACAGATGTTCCAGATAAATATTGGAAGCTATATACGTACTTTGTTTCAAAATGGGTCAATCGAGAATATCGTCAACTGTTTACTAAAAATCAGTTTCATCAAGCAATGAAACACGCCAAA* |
| ***$bla_{CTXM}$*** | *CTATGGCACCACCAACGATATCGCGGTGATCTGGCCAAAAGATCGTGCGCCGCTGATTCTGGTCACTTACTTCACCCAGCCTCAACCTAAGGCAGAAAGCCGT* |

**Table S2.** Primers used for PCR.

| ***Gene*** | ***Forward Primer (5' → 3')*** | ***Reverse Primer (5' → 3')*** |
|---|---|---|
| ***16S rRNA*** | ACTCCTACGGGAGGCAGCAG | ATTACCGCGGCTGCTGG |
| ***qnrS*** | GACGTGCTAACTTGCGTGAT | TGGCATTGTTGGAAACTTG |
| ***intI1*** | GATCGGTCGAATGCGTGT | GCCTTGATGTTACCCGAGAG |
| ***sul1*** | CGCACCGGAAACATCGCTGCAC | TGAAGTTCCGCCGCAAGGCTCG |
| ***sul2*** | TCCG**G**TGGAGGCCGGTATCTGG | CGGGAATGCCATCTGCCTTGAG |
| ***ermB*** | AAAACTTACCCGCCATACCA | TTTGGCGTGTTTCATTGCTT |
| ***$bla_{CTXM}$*** | CTATGGCACCACCAACGATA | ACGGCTTTCTGCCTTAGGTT |

**Table S3.** Mobilome number of primary reads after INTEGRALL, ISfinder and ICEberg alignment

| *Step* | ***eDNA*** | ***iDNA*** |
|---|---|---|
| *Number of INTEGRALL primary aligned* | 411 | 4'586 |
| *Number of ISfinder primary aligned* | 140 | 20'046 |
| *Number of ICEberg (ICE) primary aligned* | 77 | 9'550 |

**DEAE column showed high selectivity and efficiency towards nucleic acids from water samples**

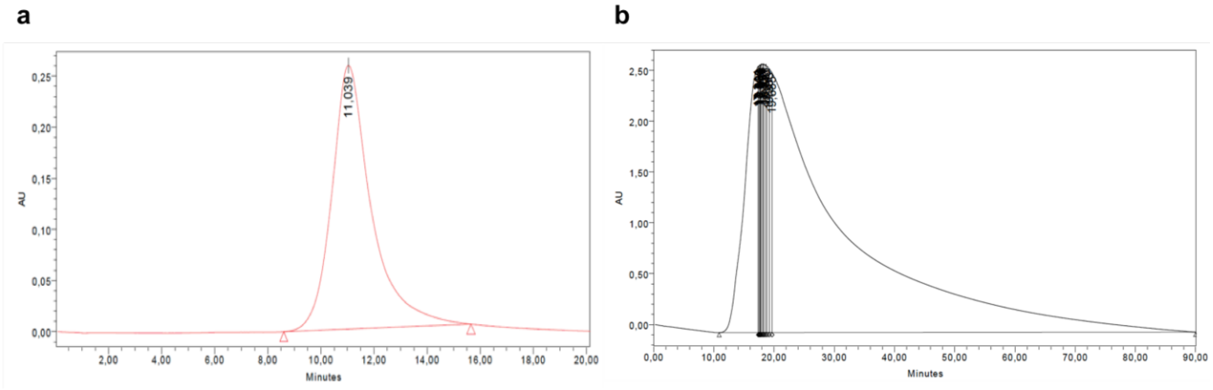


**Figure S1.** Control chromatogram (λ=260 nm) with **(a)** solution containing pure plasmid pHT01 (0.73 ng µL$^{-1}$, 10 mL) **(b)** Raw eDNA from filtered activated sludge water sample (900 mL) chromatogram (λ=260 nm).

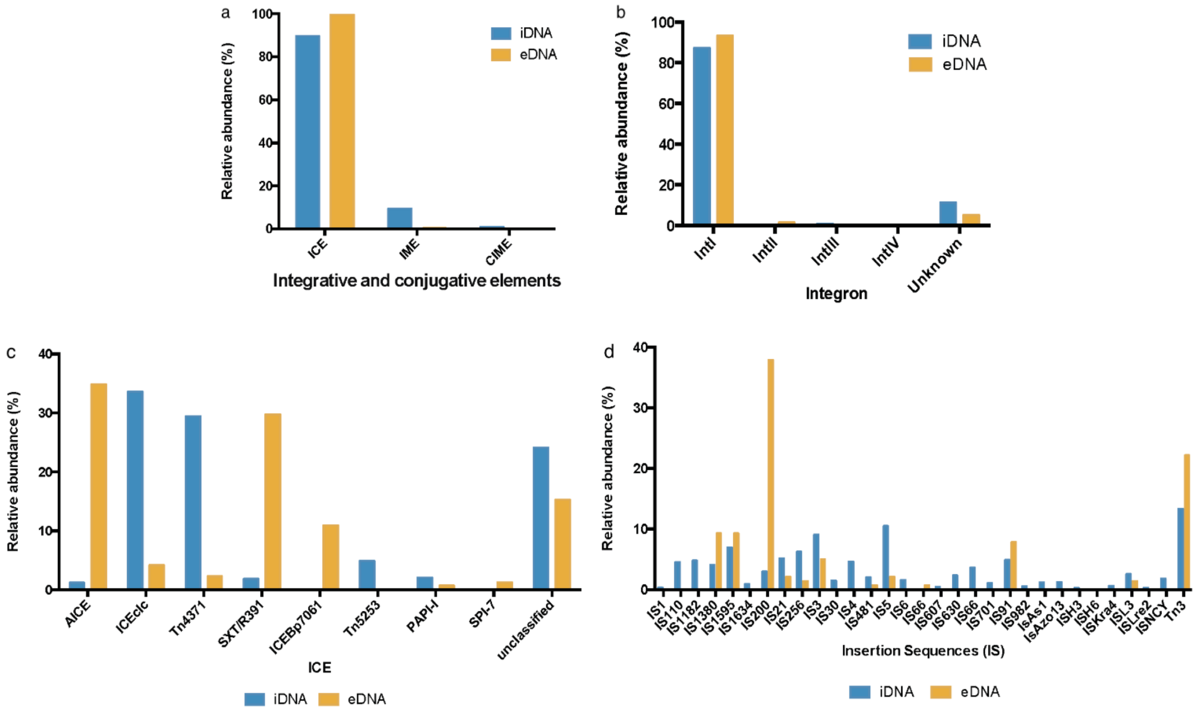


**Figure S2.** Mobilome analysis from eDNA and iDNA fractions obtained from an activated sludge sample **(a)** Integrative, conjugative (ICE), mobilizable (IME) and cis-Mobilizable elements (CIME) relative abundance. **(b)** Integrons relative abundance. **(c)** Integrative conjugative elements (ICE) relative abundance and **(d)** insertion sequences relative abundance.
